# Engulfment of viable neurons by reactive microglia in prion diseases

**DOI:** 10.1101/2024.03.06.583759

**Authors:** Natallia Makarava, Tarek Safadi, Olga Mychko, Narayan P. Pandit, Kara Molesworth, Simone Baiardi, Li Zhang, Piero Parchi, Ilia V. Baskakov

**Author notes:** Address correspondence to: Ilia V. Baskakov, Center for Biomedical Engineering and Technology, University of Maryland School of Medicine, Baltimore, 111 S. Penn St., Baltimore, MD, 21201, USA.

## Abstract

Microglia are recognized as the main cells in the central nervous system responsible for phagocytosis. During brain development, microglia eliminate excessive synapses and neurons, whereas in normal aging and neurodegenerative diseases, microglia are responsible for clearing protein aggregates and cell debris. The current study demonstrates that in prion disease, microglia effectively phagocytose prions or PrP^Sc^ during early preclinical stages. However, during the late preclinical stage, a critical shift occurs in microglial activity from PrP^Sc^ uptake to the engulfment of neurons. This change occurs before the manifestation of clinical symptoms and is followed by a rapid accumulation of total PrP^Sc^, suggesting a potential link to neuronal dysfunction and behavioral deficits. Surprisingly, the engulfed neurons do not show apoptotic markers, indicating that microglia are targeting viable neurons. Despite up to 40% of neurons being partially engulfed at the clinical stage, there is no significant neuronal loss, suggesting that many engulfment events are incomplete, terminated or protracted. This phenomenon of partial engulfment by reactive microglia is independent of the CD11b pathway, previously associated with phagocytosis of newborn neurons during neurodevelopment. The study establishes partial engulfment as a consistent occurrence across multiple prion-affected brain regions, various mouse-adapted strains, and different subtypes of sporadic Creutzfeldt-Jakob disease (sCJD) in humans. The current work describes a new phenomenon of partial engulfment of neurons by reactive microglia, shedding light on a novel aspect of neuronal-microglia interactions.

## Introduction

Microglia are recognized as the main cells in the central nervous system (CNS) responsible for phagocytosis. In early brain development, microglia are crucial in optimizing neural circuitry and eliminating excessive synapses, neurons, astrocytes, and oligodendrocytes [2, 3]. Throughout adult life, microglia defend the CNS against invading pathogens and clean up tissues from apoptotic cells and myelin debris. During normal aging and neurodegenerative diseases, microglia encounter new challenges, as they are responsible for clearing protein aggregates and cell debris, while simultaneously participating in repair and regeneration processes. As a response to ongoing changes in brain homeostasis and persistent exposure to protein aggregates, microglia acquire various reactive phenotypes, some of which are characterized by upregulated phagocytic pathways [1, 4–6]. It has been proposed that the upregulation of the phagocytosis under conditions of chronic neuroinflammation leads to a phagocytosis of viable neurons by microgia, contributing to neurodegeneration [7]. Indeed, it appears that the phagocytic pathways responsible for pruning of synapses and neurons during neurodevelopment also contribute to synaptotoxicity in neurodegenerative diseases and aging [7–10]. However, most of the evidence that microglia can phagocytose viable neurons were collected *in vitro* using cultured cells, whereas detecting engulfment of viable neurons in actual neurodegenerative disease has been difficult [11–13].

Phagocytosis comprises several sequential steps, encompassing the recognition, engulfment, and degradation of a target [14]. When microglia encounter a target, they extend filopodia, forming a phagocytic cup that engulfs the target and gives rise to phagosomes [14]. Subsequently, phagosomes undergo maturation by merging with endosomes and lysosomes, forming phagolysosomes, where the phagocytosed material is digested. *In vivo*, the entire process of microglia-mediated phagocytosis of an individual cell is completed in less than two hours [15].

Prion diseases, also known as Transmissible Spongiform Encephalopathies, represent a group of transmissible neurodegenerative disorders affecting both humans and animals [16]. These conditions currently lack any effective treatment, and their invariably fatal outcome is well-established. Prion diseases are instigated by prions or PrP^Sc^, which is the misfolded, aggregated form of a cellular sialoglycoprotein known as the prion protein or PrP^C^ [17]. The pathogenic mechanism involves the replication and dissemination of prions throughout the CNS, achieved by recruiting and converting host-expressed PrP^C^ molecules into misfolded, β-sheet-rich PrP^Sc^ states [18]. In individuals or animals infected with prions, the neurodegeneration is attributed to PrP^Sc^ accumulation, which exerts a toxic effect on neurons [19–25].

Over the years, the idea that microglia constitute the primary host defense against prions has gained strong experimental support [26–28]. Indeed, the ablation of microglia, either before prion infection or during the initial stages of the disease, significantly accelerated disease progression [29–32]. Furthermore, the knockout of MFGE8, a factor secreted by microglia that mediates the phagocytosis of apoptotic bodies, resulted in a 40-day acceleration of prion pathogenesis and elevated levels of PrP^Sc^ [33]. Contrary to the notion that microglia are protective, partial inhibition of microglia proliferation and reactivity at the preclinical stage (98 to 126 days post-infection with the ME7 strain) delayed the onset of behavioral signs and extended survival by 26 days [34]. Moreover, the inhibition of microglia activation through the administration of an immunosuppressant just before or at the time of disease onset suppressed reactive gliosis and prolonged the survival of humanized mice infected with human prions [35]. In response to prion infection, microglia was shown to upregulate IFN-1 (interferon type 1), which activates phagolysosomal pathways [36]. Surprisingly, the knockout of the *IFNAR1* gene, which encodes a receptor for IFN-1, ameliorated clinical signs, improved neuronal survival along with synaptic density, and prolonged survival in ME7-infected mice [36].

In our present study, we demonstrate that microglia efficiently phagocytose PrP^Sc^, starting in the early preclinical stage, thereby helping to maintain the total PrP^Sc^ at low levels. However, during the late preclinical stage, there is a shift in the target of microglial activity from PrP^Sc^ uptake to the engulfment of neurons. This transition in microglial phagocytic behavior is followed by a rapid accumulation of total PrP^Sc^. Notably, microglia that encircle neurons exhibit hypertrophic lysosomal compartments, suggesting their readiness for a phagocytic engagement. Importantly, neuronal engulfment begins prior to clinical onset, implicating its potential contribution to neuronal dysfunction and behavioral deficits. Surprisingly, neurons undergoing engulfment lack apoptotic markers, indicating that reactive microglia are targeting viable neurons. Despite the presence of up to 40% partially engulfed neurons at clinical stages, the absence of neuronal loss suggests that the majority of engulfing events are incomplete or arrested. Indeed, the vast majority of neurons are only partially engulfed, suggesting that engulfment is often stalled or terminated. Neuronal engulfment is found to be independent of the CD11b pathway, which was previously identified as responsible for the phagocytosis of newborn neurons during neurodevelopment. The current work describes a new phenomenon of partial encircling of neurons by reactive microglia that resembles engulfment. Partial engulfment was consistently observed across multiple prion-affected brain regions, four mouse-adapted strains examined here, and various subtypes of sCJD in humans.

## Materials and Methods

### Animals

10% (w/v) SSLOW, RML, 22L or ME7 brain homogenates (BH) for inoculations were prepared in PBS, pH 7.4, using glass/Teflon homogenizers attached to a cordless 12 V compact drill as previously described [37]. SSLOW, RML, 22L and ME7 brain-derived materials were inoculated as 1% or 10% BH via i.p. or i.c. route. Immediately before inoculation, each inoculum was further dispersed by 30 sec indirect sonication at ∼200 watts in a microplate horn of a sonicator (Qsonica, Newtown, CT). Each mouse received 20 μl of inoculum i.c. or 200 μl i.p. under 3% isoflurane anesthesia. After inoculation, animals were observed regularly for signs of neurological disorders. Clinical signs included clasping hind legs, difficulty walking, abnormal gait, nesting problems, and weight loss. The animals were deemed symptomatic when at least two clinical signs were consistently observed. The mice were euthanized when they were unable to rear and/or lost 20% of their weight. CD11b^−/−^ mice with a global knockout of CD11b used in the current work were on the C57Bl/6J background, and are described elsewhere [38]. Normal control groups were age-matched non-inoculated mice. For caspase-3 immunostaining positive control, mice were subjected to transient middle cerebral artery occlusion (MCAO) as previously described [39]. 5XFAD mice were bred and housed as previously described [39].

### Elevated plus maze (EPM) test and scoring of clinical signs of prion disease

Mice were tested once per week starting at the preclinical stage. In each session, a mouse was placed at the center of a plus maze and given 5 minutes to explore the maze. All movements were captured using video recording, and total timing spent in open arms, closed arms or the center of the maze was quantified using ANY-maze software (version 6.33). After the first training session, mice naturally acquire a strong preference for the closed arms, i.e. avoiding open arms during the sessions that follow the training session until the clinical onset of the disease. The clinical onset was defined at a time point when mice lose their preference for the closed arms.

### Antibodies

Primary antibodies used for immunofluorescence, immunohistochemistry and immunoblotting were as follows: rabbit polyclonal anti-IBA1 (#013-27691, FUJIFILM Wako Chemicals USA; Richmond, VA); goat polyclonal anti-IBA1 (#NB100-1028, Novus, Centennial, CO); chicken polyclonal anti-MAP2 (#NB300-213, Novus, Centennial CO); mouse monoclonal anti-NeuN, clone A60 (#MAB377, Millipore Sigma, Burlington, MA); rat monoclonal anti-CD68, clone FA-11 (#MCA1957, BioRad, Hercules, CA); mouse monoclonal anti-PV, clone PARV-19 (#MAB1572, Millipore Sigma, Burlington, MA); rabbit polyclonal anti-Caspase 3, active (cleaved) form (#AB3623, Millipore Sigma, Burlington, MA); mouse monoclonal anti-prion protein, clone SAF-84 (#189775, Cayman, Ann Arbor, MI); rabbit monoclonal anti-prion protein, clone 3D17 (#ZRB1268, Millipore Sigma, Burlington, MA); rabbit monoclonal anti-GFAP, clone D1F4Q (#12389, Cell Signaling Technology, Beverly, MA); chicken polyclonal anti-GFAP (#AB5541, Millipore Sigma, Burlington, MA); mouse monoclonal anti-β-amyloid, clone 6E10 (#803004, BioLegend, San Diego, CA); rat monoclonal anti-LAMP1, clone 1D4B (#121601, BioLegend, San Diego, CA); rabbit polyclonal anti-CD11b (#ab128797, Abcam, Boston, MA); mouse monoclonal anti-Tubulin β3, clone TUJ1 (#801213, BioLegend, San Diego, CA); rat monoclonal anti-Gal3, clone M3/38 (#sc-23938, Santa Cruz, Dallas, TX); mouse monoclonal anti-β-actin, clone AC-15 (#A5441, Sigma-Aldrich, Saint Luis, MO). The secondary antibodies for immunofluorescence were Alexa Fluor 488-, 546-, and 647-labeled (ThermoFisher Scientific, Waltham, MA).

### Western blot and densitometry

For Western blot, 10% (w/v) brain homogenates (BH) were prepared in RIPA Lysis Buffer (Millipore Sigma, St. Louis, MO) using glass/Teflon homogenizers attached to a cordless 12V compact drill as previously described [37]. To detect PrP^Sc^, 10% BHs were mixed with an equal volume of 4% sarcosyl in PBS, supplemented with 50 mM Tris, pH 7.5, and digested with 20 μg/ml PK (New England BioLabs, Ipswich, MA) for 30 min at 37°C with 1000 rpm shaking, using an Eppendorf thermomixer. PK digestion was stopped by adding SDS sample buffer and heating the samples for 10 min in a boiling water bath. Samples were loaded onto NuPAGE 12% Bis-Tris gels (ThermoFisher Scientific, Waltham, MA), transferred to a PVDF membrane, and probed with the indicated antibodies according to standard protocols. Western blot visualization and densitometry were performed with Immobilon Forte Western HRP substrate or SuperSignal West Pico PLUS Stable Peroxidase and Luminol/Enhancer, using iBright 1500 imaging system and software (ThermoFisher Scientific, Waltham, MA).

### Immunofluorescence and DAB staining of mouse brains

Formalin-fixed brains (sagittal or coronal 3 mm slices) were treated for 1 hour in 96% formic acid before being embedded in paraffin using standard procedures. 4 μm sections produced with Leica RM2235 microtome (Leica Biosystems, Buffalo Grove, IL) were mounted on Superfrost Plus Microscope slides (#22-037-246, Fisher Scientific, Hampton, NH) and processed for immunohistochemistry according to standard protocols. To expose epitopes, slides were subjected to 20 min of hydrated autoclaving at 121° C in Citrate buffer, pH6.0, antigen retriever (#C9999, Sigma-Aldrich, Saint Luis, MO). For the detection of disease-associated PrP, an additional 3 min treatment in concentrated formic acid was applied.

For immunofluorescence, an Autofluorescence Eliminator Reagent (Sigma-Aldrich, St. Louis, MO) and Signal Enhancer (ThermoFisher, Waltham, MA) were used on slides according to the original protocols to reduce background fluorescence. Epifluorescent images were collected using an inverted microscope Nikon Eclipse TE2000-U (Nikon Instruments Inc, Melville, NY) equipped with an illumination system X-cite 120 (EXFO Photonics Solutions Inc., Exton, PA) and a cooled 12-bit CoolSnap HQ CCD camera (Photometrics, Tucson, AZ). Images were processed using FIJI software (ImageJ 2.14.0, National Institute of Health, Bethesda, MD, United States).

Detection with 3,3′-diaminobenzidine (DAB) was performed using horseradish peroxidase labeled secondary antibodies and DAB Quanto chromogen and substrate (VWR, Radnor, PA).

### Immunohistochemistry of sCJD brains

Immunohistochemistry of 57 sCJD brains, including 42 from the three most common disease subtypes (MM1, VV2 and MV2K) was performed as described before [40]. Briefly, 7 μm sections from formalin-fixed and paraffin embedded tissue blocks were stained using a monoclonal antibody against the b-chain of human HLA-DR, –DQ, –DP (clone CR3/43, 1:400; Agilent Dako). The anti-HLA antibody was chosen after carrying out preliminary comparative experiments with other antibodies known to detect activated microglia, such as CD68 or Iba1, based on accuracy and resolution. Antigen retrieval was achieved by microwaving the sections in sodium citrate buffer (0.01 M, pH 6.0, 15 min). Slides were then loaded into an Aperio ScanScope XT (Aperio, Leica Biosystems, USA) and scanned at 20x magnification (1 μm/pixel) via the semi-automated method. Slides were checked for image quality using an Aperio ‘quality factor >90’ and visual inspection, and annotated using the ImageScope v12 software (Leica Biosystems, USA).

### TUNEL assay

Click-IT Plus TUNEL Assay (#C10617, ThermoFisher, Waltham, MA) was performed on brain sections prepared without formic acid treatment. The TUNEL assay protocol recommended by the manufacturer was adapted for staining of free-floating sections within a 24-well plate. Following the TUNEL reaction, brain sections were immunostained for IBA1 to allow the visualization of microglia.

### Confocal microscopy and 3D image reconstruction

Confocal images were acquired using a Leica TCS SP8 microscope using the 40x/1.30 or 63x/1.40 oil immersion objective lenses with laser lines 405, 488, 552, and 638 when needed. Image size was 1024 x 1024 pixels at a 400 Hz scan speed. Z stack thickness ranged from 6.2 to10.45 μm taken at the system-optimized number of steps. Images were processed using the LAS X and FIJI software.

### Quantification of engulfment and neuronal count

To estimate the percentage of neurons engulfed by microglia (Fig. 2F, 4B, and 9F), 3 – 5 brains from each animal group were co-immunostained for MAP2 and IBA1, and multiple images from cortices of each brain were collected under 60x objective, as noted in the Figure Legends. Using FIJI software, images obtained from green and red channels were subjected to an automated threshold, and for each MAP2^+^ neuron, an area of overlap between signals from green (MAP2) and red (IBA1) channels was recorded. Based on the calculations for normal brains, 20-pixel overlap was chosen as a cut-off for the detection of engulfment events. Neurons were counted as undergoing engulfment, if they had over 20 pixels of MAP2 signal overlapped by IBA1 signal. The total number of identified MAP2^+^ cells was recorded as a neuronal count.

**Figure 1.**
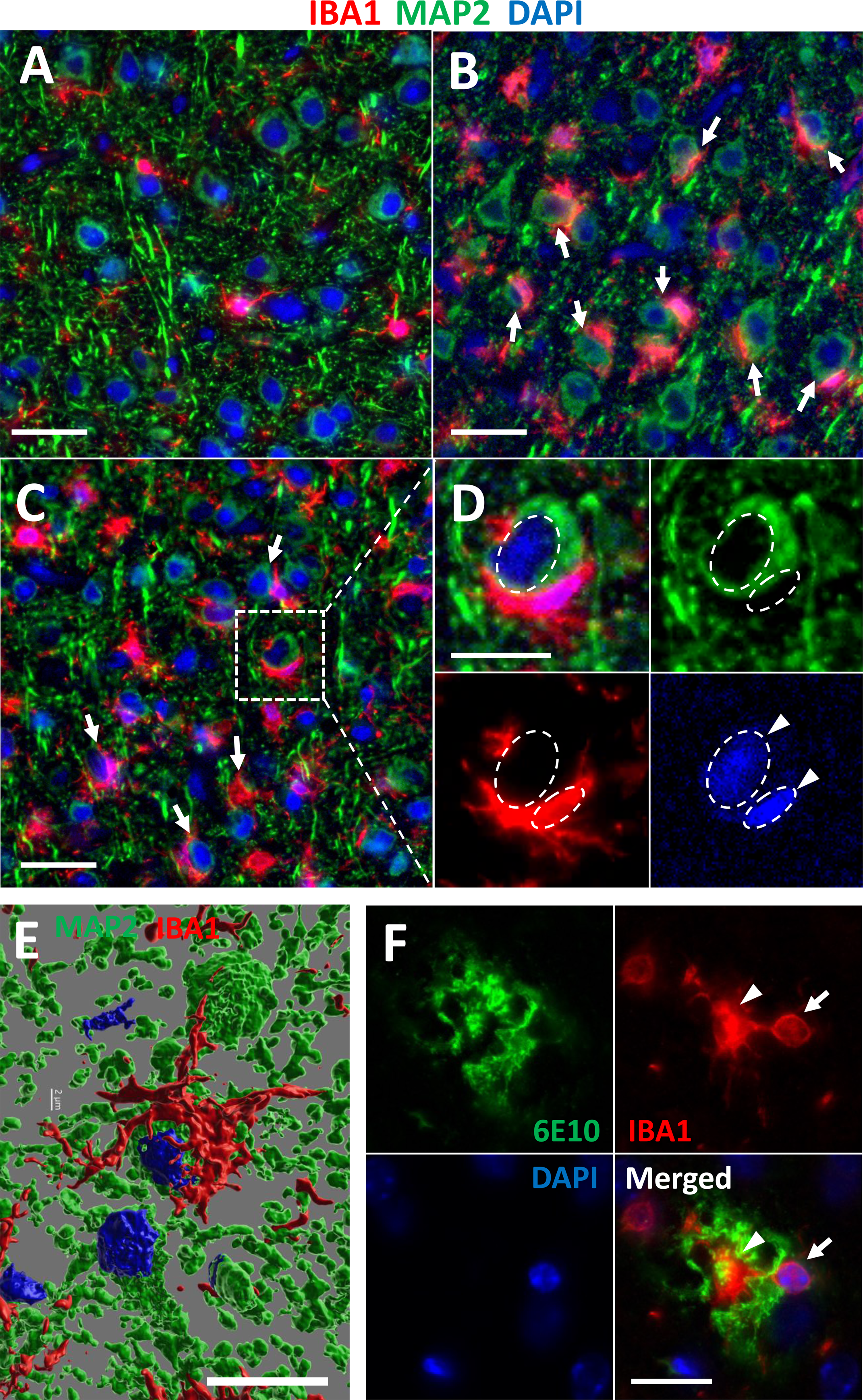
Partial engulfment of neurons by reactive microglia in prion-infected mice. (**A-E**) Coimmunostaining of terminally ill C57Bl/6J mice infected with SSLOW via i.c. (**B**) or i.p. routes (**C-E**), and mock controls (**A**) using anti-IBA1 (a marker of microglia, red) and anti-MAP2 (a neuronal marker, green) antibodies. Arrows point at partially engulfed neurons. (**D**) Enlarged image of engulfment showing cup-shaped microglial soma. Arrowheads and dashed circles mark neuronal and microglial nuclei located in close proximity. (**E**) 3D confocal reconstruction of partial engulfment of a neuron by reactive microglia. (**F**) Co-immunostaining of 10-month-old 5XFAD mice using anti-IBA1 (red) and 6E10 antibody (green) that stains Aβ amyloid plaques. The arrow points at microglia soma, and the arrowhead points at thee phagocytic cup. Scale bars = 20 µm in A-C and F, 10 µm in D and E.

**Figure 2.**
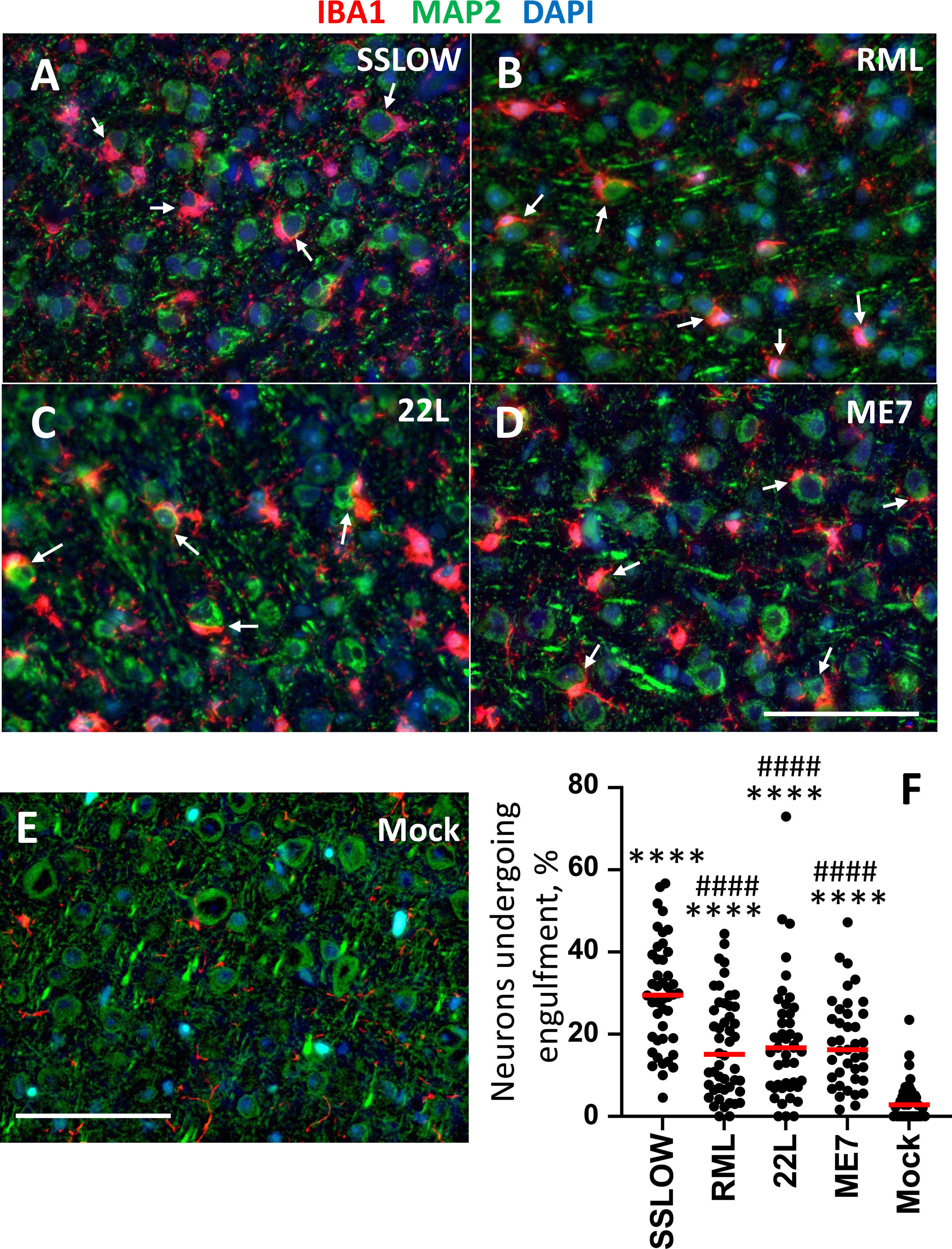
Engulfment of neurons is a common property among prion strains. (**A-D**) Representative images of partial engulfment of neurons by reactive microglia in cortex of C57Bl/6J mice infected with SSLOW (**A**), RML (**B**), 22L (**C**), and ME7 (**D**) via i.p. and stained using anti-IBA1 (red) and anti-MAP2 (green) antibodies. Arrows point at neurons undergoing engulfment. (**E**) IBA1/MAP2 immunostaining of mock-inoculated age-matched normal brains. (**F**) Quantification of the percentage of neurons that are undergoing engulfment by microglia in cortexes of mice infected i.p. with SSLOW, RML, 22L or ME7, and mock control mice. **** p<0.0001 – statistical significance vs mock; #### p<0.0001 – statistical significance vs SSLOW by non-parametric Mann-Whitney test (N=4 individual animals per group, n=39-48 fields of view per group). Scale bars = 50 µm.

**Figure 3.**
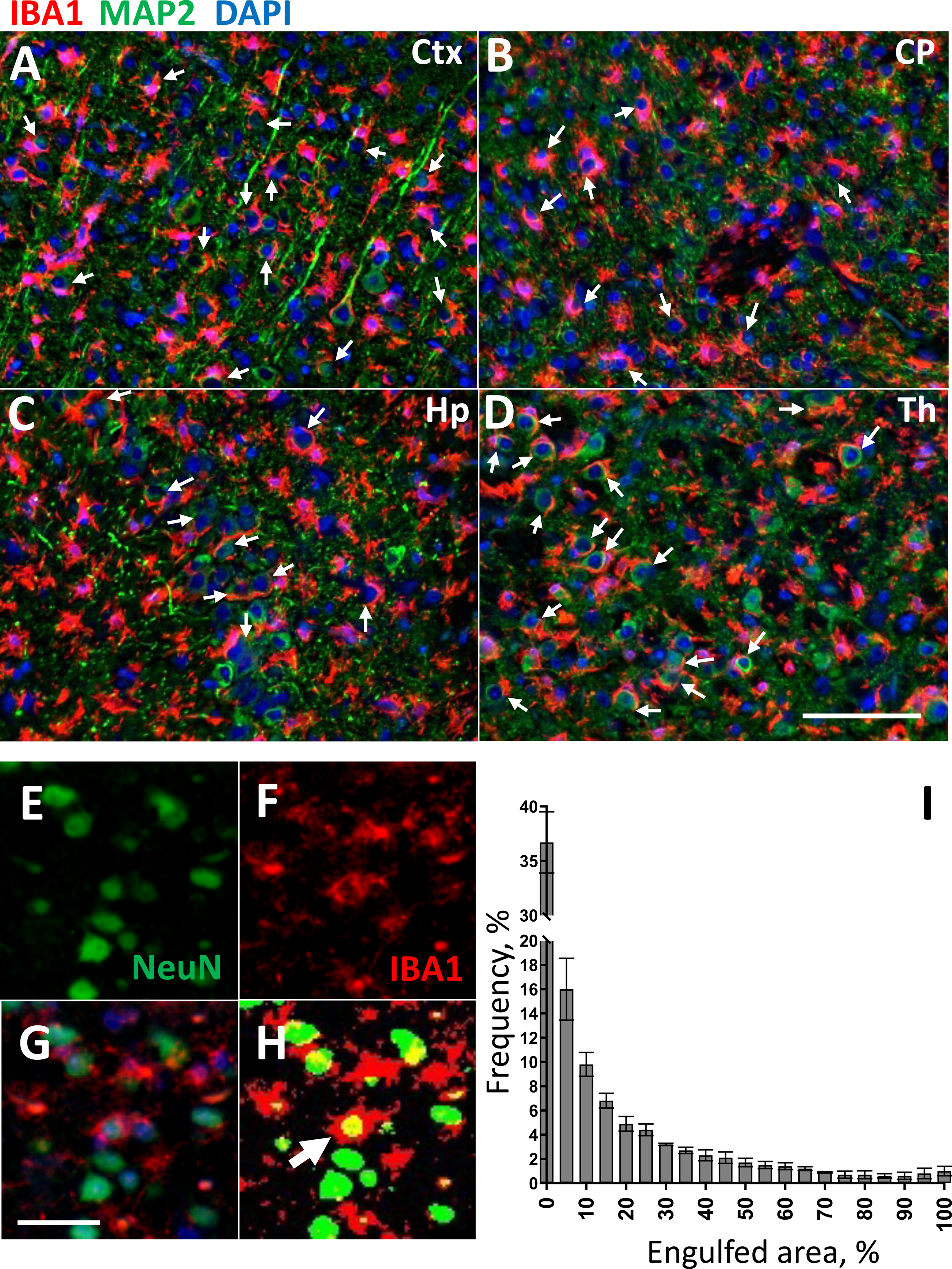
The vast majority of neurons are only partially engulfed. (**A-D**) Engulfment of neurons in cerebral cortex (**A**), caudate/putamen (**B**), hippocampus (**C**), and thalamus (**D**) of C57Bl/6J mice infected with SSLOW via i.c. route at the terminal stage of the disease. Staining was performed using anti-IBA1 (red) and anti-MAP2 (green) antibodies. Arrows point to examples of neuronal engulfment. (**E-I**) Quantification of the neuronal area engulfed by microglia. With epifluorescence microscopy, light penetrates the full depth of a cell, thus an overlap of fluorescence signal from neuronal (NeuN, green, **E**) and microglial (IBA1, red, **G**) markers is observed when a neuronal body is undergoing engulfment by a microglial cell. The percentage of neuronal area engulfed by microglia is estimated for individual neurons as a fraction of NeuN signal overlapped with IBA1 signal (**G**). (**H**) Merged thresholds of images from IBA1 and NeuN channels. The arrow points to a very rare event when the NeuN signal is overlapped with the IBA1 signal close to 100%, which is counted as a complete engulfment. (**I**) Frequency distribution of the engulfed areas of the individual neurons, as quantified for the cortex of SSLOW-infected mice. N=3 individual animals, n=1169, 1422 and 1551 engulfing events for individual animals. Scale bars = 100 µm for A-D and 20 µm for E-H.

**Figure 4.**
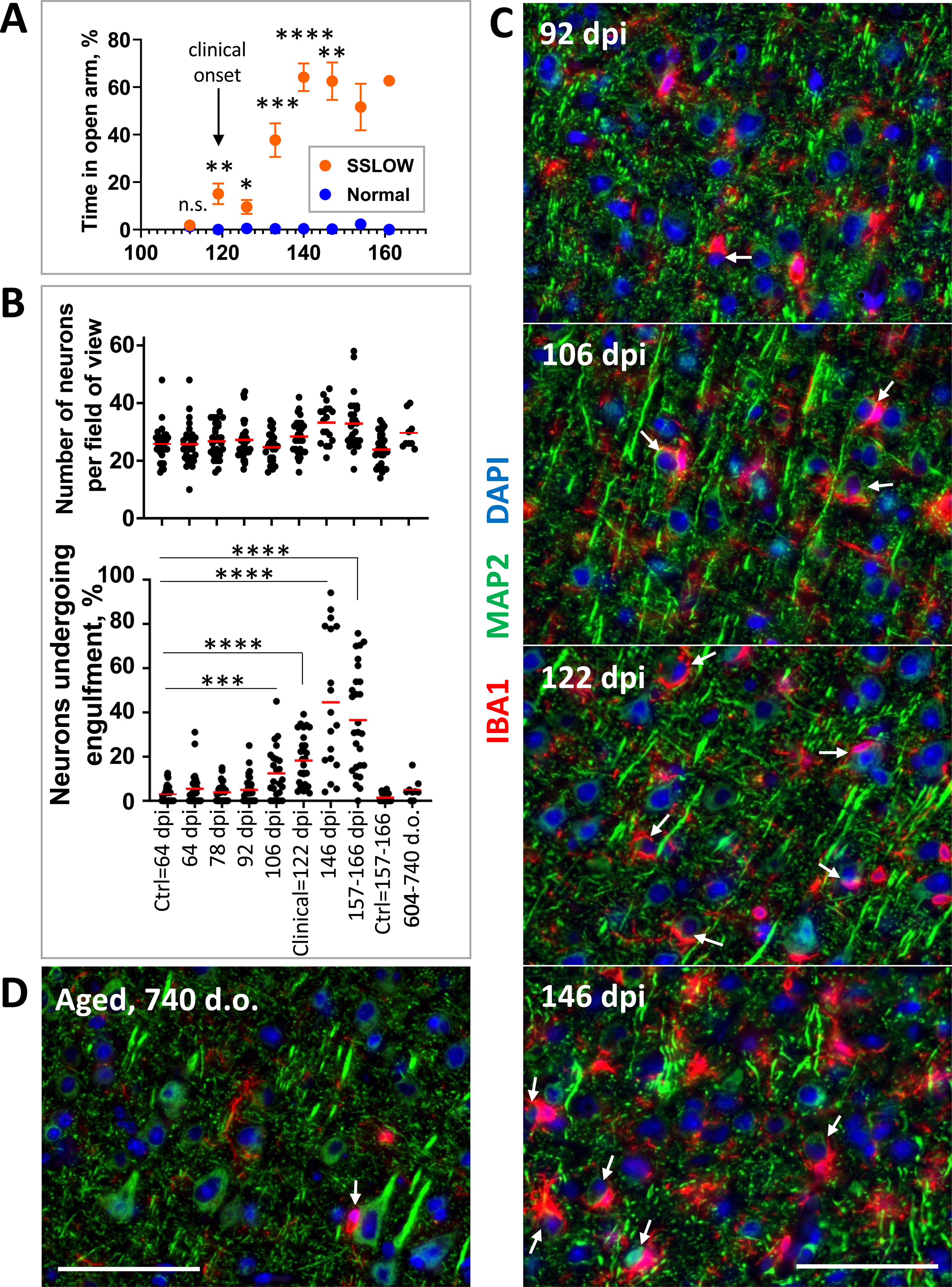
Time course of the engulfment. C57Bl/6J mice were infected i.p. with SSLOW. (**A**) The clinical onset of the disease was established using the elevated plus maze (EPM) test. Mice were subjected to EPM sessions once per week starting from the preclinical stage. After the first training sessions (not shown), mice naturally acquire a strong preference for the closed arms, i.e. avoiding open arms. The clinical onset was defined as a time point, when the time spent on open arms consistently increased in comparison to the normal age-matched group (shown by arrow). N=5 for normal; N=14 for SSLOW until 119 dpi; * p<0.05, ** p<0.01, *** p<0.001, **** p<0.0001 by Tukey’s multiple comparison test. (**B**) Change in the total number of MAP2^+^ neurons (upper plot) and the percentage of MAP2^+^ neurons undergoing engulfment (lower plot) during preclinical and clinical stages of the disease in mice inoculated i.p. with SSLOW. N=3-5 individual animals for each time point, n=18-32 fields of view (except for 604-740 dpi, where n=9 fields of view). p<0.001, **** p<0.0001 by Mann-Whitney test. (**C**) Representative images of partial engulfment of neurons by microglia in the cerebral cortex of C57Bl/6J mice infected i.p. with SSLOW and collected at 92 dpi, 106 dpi, 122 dpi (disease onset), and 146 dpi. (**D**) Representative image of aged brains (740 days old). Brain sections were stained using anti-IBA1 (red) and anti-MAP2 (green) antibodies. Arrows point at examples of neurons undergoing engulfment. Scale bars = 50 µm.

To estimate the percentage of PV^+^ neurons undergoing engulfment by microglia, brains were co-immunostained with anti-PV and anti-IBA1 antibodies, and multiple images from cortices of each brain were collected under 20x objective. Images from red (PV) and green (IBA1) channels were thresholded, converted to binary, and PV^+^ neurons were identified as regions of interest (ROIs) using the Analyze Particles function of FIJI software. The same ROIs were applied to corresponding images from the IBA1 channel, and the area of IBA1 signal overlap with each PV^+^ neuron was recorded. An area of ≥20 pxls was counted as engulfment. The total number of identified PV^+^ neurons in each image was also recorded.

To assess the frequency of complete engulfment events, 3 brains of C57Bl/6J mice infected with SSLOW via i.c. route and collected at thee terminal stage of the disease were co-immunostained for NeuN and IBA1. The images were collected from cortices of each brain under 20x objective. The neurons were identified using the Analyze Particles function of FIJI software as above. The percent of NeuN signal area covered by IBA1 signal was recorded for each neuron. The data were presented as a histogram of frequencies of neurons engulfed to a different extent, from 0% to 100%.

### Quantification of cCasp3^+^ cells

To compare the number of cCasp3^+^ neurons and microglia cells, 2 brains of terminal SSLOW-infected C57Bl/6J mice and 2 brains of normally aged (607, 740 d.o.) C57Bl/6J mice were co-immunostained for cCasp3 and NeuN or IBA1, and images from the cortices of each brain were collected under 20x objective. The images from each channel were subjected to a set threshold, converted to binary, and neurons or microglia cells were identified as regions of interest (ROIs) using the Analyze Particles function of FIJI software. The same ROIs were applied to the corresponding cCasp3 channel, and the presence of cCasp3 signal in each identified cell was measured. The data were presented as a percent of cells containing cCasp3 puncta.

### Colocalization of PrP signal with different cell types

To quantify colocalization of PrP signal with different cell types, 3 brains of terminal SSLOW-infected C57Bl/6J mice were co-immunostained for PrP (3D17) and cell type markers: MAP2 for neurons, GFAP for astrocytes, and IBA1 for microglia. The images from each channel were subjected to a set threshold, and an area of overlap between PrP signal and MAP2, GFAP or IBA1 was detected using the Image Calculator function of FIJI software. Total 3D17 signal intensity and 3D17 signal intensity in the areas of overlap were measured. The data were presented as a percent of PrP signal intensities colocalizing with neurons, astrocytes or microglia.

### Quantification of PrP^Sc+^ microglia

To estimate the number of PrP^Sc+^ microglia cells, brains of C57Bl/6J non-infected control mice and mice infected with SSLOW via i.p. route, collected at various time points, were co-immunostained with anti-PrP 3D17 and IBA1 antibodies. Images from thee cortices of each brain were collected under 60x objective. The images from each channel were subjected to a set threshold, converted to binary, and microglia cells were identified as regions of interest (ROIs) using the Analyze Particles function of FIJI software. The same ROIs were applied to the corresponding 3D17 channel, and the presence of PrP^Sc^ signal in each identified microglia cell was measured. The number of microglia cells identified in each image was also recorded. The data were presented as a percent of microglia cells containing PrP^Sc^.

### Quantification of LAMP^+^ in microglia

IBA and LAMP1 images collected with confocal microscopy were merged, and background was subtracted. Then, single microglial cells were selected and opened as separate images. The images were split into channels and thresholded using the Yen method. The area and mean grey value of each microglia cell were measured, and the area of LAMP1 inside microglia was determined using the Image Calculator function with the AND operation. Mean grey value of LAMP1 inside microglia was determined by overlaying LAMP1 binary selection from previous step onto the original image and measuring intensity.

### Analysis of gene expression

After euthanasia by CO2 asphyxiation, brains were immediately extracted and kept ice-cold during dissection. Brains were sliced using a rodent brain slicer matrix (Zivic Instruments, Pittsburg, PA). 2 mm central coronal sections of each brain were used to collect thalamus and cortex. Allen Brain Atlas digital portal (http://mouse.brain-map.org/static/atlas) was used as a reference. RNA isolation was performed as described before [41]. RNA samples were processed by the Institute for Genome Science at the University of Maryland School of Medicine using the nCounter custom-designed NanoString gene panel (NanoString Technologies, Seattle, WA). Only samples with an RNA integrity number RIN > 7.2 were used for NanoString analysis. All data passed quality control, with no imaging, binding, positive control, or CodeSet content normalization flags. The analysis of data was performed using nSolver Analysis Software 4.0. Ten house-keeping genes ( *Xpnpep1, Lars, Tbp, Mto1, Csnk2a2, CCdc127, Fam104a, Aars, Tada2b, Cnot10*) were used for normalization of gene expressions.

### Statistics

Statistical analyses and plotting of the data were performed using GraphPad Prism software, versions 8.4.2 – 10.1.1 for Windows (GraphPad Software, San Diego, California USA) or Excel 2016 – version 2302, as detailed in Figure Legends.

### Study approval

The study was carried out in strict accordance with the recommendations in the Guide for the Care and Use of Laboratory Animals of the National Institutes of Health. The animal protocol was approved by the Institutional Animal Care and Use Committee of the University of Maryland, Baltimore (Assurance Number: A32000-01; Permit Number: 0215002). Human brain tissues have been collected under the Italian National Surveillance program for CJD and related disorders, and their use for research was approved by written informed consent of patients during life or their next of kin after death.

## Results

### In prion-infected mice, reactive microglia engulf neurons

Co-immunostaining of clinically ill C57Bl/6J mice infected with the mouse-adapted strain SSLOW, administered via intraperitoneal (i.p.) or intracranial (i.c.) routes, revealed a substantial population of neurons in close contact with the soma of reactive microglia (Fig. 1A-D). In normal, non-infected cortices, such contacts between neurons and microglial soma were infrequent (Fig. 1A). Three-dimensional reconstruction of confocal imaging revealed a partial encircling of neurons by reactive microglia in prion-infected mice (Fig. 1E). The vast majority of neurons in contact with microglia were only partially encircled (Fig. S1A, B).

Microglia typically execute phagocytosis by forming pseudopodia with a phagocytic cup [14, 42, 43]. This cup engulfs the target, which may include synapses, apoptotic cells, or protein aggregates. Full engulfment of a target by the phagocytic cup results in the formation of phagosomes within pseudopodia. Consistent with this mechanism, pseudopodia with phagocytic cups were observed in adult 5XFAD mice, where microglia engulfed Aβ aggregates (Fig. 1F) or neurons (Fig. S1C). In contrast to the traditional phagocytic mechanism involving pseudopodia and a phagocytic cup, in prion-infected mice, microglial soma surrounded neurons (Fig. 1D, S1B). The proximity of microglial and neuronal nuclei supports the notion that neurons were encircled by microglial somas, which often exhibited a characteristic cup-shaped appearance (Fig. 1D, S1B) rather than by extended pseudopodia. Engulfment of individual neurons by microglia in prion-infected brains occurred at a one-to-one ratio (Fig. 1B-E, S1B).

### Neuronal engulfment is a common phenomenon observed across prion strains

C57Bl/6J mice, infected with four prion strains (SSLOW, RML, 22L, and ME7), exhibited partial engulfment of neurons by reactive microglia (Fig. 2A-D, F). Notably, by the terminal stage of the disease, SSLOW mice showed the highest percentage of partially encircled neurons. Among the four strains, SSLOW characterized by the shortest incubation time, displayed the most pronounced interaction between microglial soma and neurons (Fig. 2A,F). In age-matched mock controls, close contact between microglial soma and neurons was infrequent (Fig. 2E,F).

Mouse-adapted prion strains employed here display distinctive strain-specific cell tropism. For instance, 22L PrP^Sc^ was predominantly associated with astrocytes, while ME7 PrP^Sc^ exhibited a prevalent localization with neurons [44]. In the case of RML, the preference of PrP^Sc^ for astrocytes versus neurons is influenced by a brain region [44]. Despite these strain-specific cell tropisms, the observation of neuronal engulfment across all strains, coupled with comparable percentages of engulfment among ME7 (mean 17.9%), 22L (mean 18.1%), and RML (mean 17.1%) strains, irrespective of their cell tropism, implies that the engulfment is not solely driven by PrP^Sc^ association with neurons (Fig. 2F).

### The vast majority of neurons exhibit only partial engulfment

Reactive microglia engulfed neurons across all brain regions affected by prions, including cortex, caudate/putamen (or striatum), hippocampus, and thalamus (Fig. 3A-D). In all these regions, most of the engulfing events were incomplete, in contrast to the relatively abundant full engulfment of cells reported during neurodevelopment, such as the phagocytosis of neuronal progenitor cells or oligodendrocytes [15, 45].

To estimate the degree of engulfment for individual neuronal cells, we calculated the area of NeuN signal covered by IBA1 signal (Fig. 3E-I). In the cerebral cortex of SSLOW-infected mice analyzed at the terminal stage, approximately 36% of neurons showed no contact with microglia, while the remaining neurons exhibited varying degrees of overlap with the IBA1 signal. Intriguingly, the complete overlap of the entire neuronal signal with the IBA1 signal was observed in only a very small percentage of neurons (less than 1%). Consequently, the majority of neurons were found to be only partially encirculed. This finding raises the possibility that in the majority of cases, the process of engulfment is arrested or terminated before the full engulfment of an entire neuronal soma.

### The engulfment of neurons by microglia does not lead to a reduction in the total number of neurons in a mouse brain

Previous studies have shown that the phagocytic clearance of a cell by microglia in a mouse brain takes 25 minutes to 2 hours [15, 45]. In the terminal stage of prion disease, as few as 15.1% of neurons (for RML) were observed to undergo engulfment (Fig. 2F). If 15% or more of the neuronal population in prion-infected animals are phagocytosed every 2 hours, the entire population would be cleared in less than 12 hours. Consequently, we aimed to establish a timeline for neuronal loss in prion-infected brains.

In C57Bl/6J mice challenged with SSLOW via i.p. route, clinical onset occurred around 120 days post inoculation (dpi) (Fig. 4A). Mice were euthanized at 157-166 dpi, when they showed 20% weight loss along with severe motor impairment and behavior deficits (Fig. 4A). Neuronal quantification and the percentage of neurons under engulfment were assessed in cortices at regular time points, starting at 64 dpi. At 64 dpi, 78 dpi, and 92 dpi, the percentage of neurons under engulfment was comparable to the control group, matched in age to the 64 dpi group (Fig. 4B). However, a statistically significant increase in the percentage of neurons undergoing engulfment was noted at 106 dpi (mean 12.4% vs 3.0% in control), two weeks prior to clinical onset (Fig. 4B). The percentage of neurons undergoing engulfment appeared to rise with the disease progression, peaking at 146 dpi (mean 44.5%) (Fig. 4B). Thus, despite a significant number of MAP2^+^ neurons being partially engulfed at the clinical stage of the disease, the total number of MAP2^+^ cells in the cortex did not decrease with the disease progression, suggesting that engulfment does not result in neuronal death (Fig. 4B). Since mice must be euthanized at 20% weight loss, defined as the endpoint per requirements of the animal care committee, it remains unknown whether significant neuronal loss would occur after this point and upon reaching the actual terminal stage of the disease.

In a cohort of aged C57Bl/6J mice (604-704 days old), the number of neurons undergoing engulfment was comparable to that in two control groups, matched in age either to the 64 dpi or 157-166 dpi groups (Fig. 4B). This observation suggests that the phenomenon of partial engulfment is not prominently associated with normal aging.

The analysis of gene expression in SSLOW-inoculated mice revealed that, despite the absence of a decline in the neuronal population, neuron-specific genes were downregulated. Notably, genes associated with ligand-gated ionic channels related to memory and learning (*Gabrg1, Grin1, Grin2b, Grm2*) and synaptic neurotransmission (*Snap25, Syn2, Syp*) exhibited reduced expression levels (Fig. S2).

In summary, the above findings establish that neuronal engulfment events were on the rise prior to the clinical onset of the disease. Surprisingly, neuronal engulfment was not accompanied by neuronal loss, suggesting that the process of engulfing was arrested or stagnant. These findings explain why in a vast majority of cases, the engulfment was only partial. Nevertheless, the engulfment appears to result in neuronal dysfunction in the absence of neuronal death and may even contribute to behavioral deficits.

### The engulfment is not selective toward the sub-population of the most vulnerable PV^+^ neurons

In previous studies, GABAergic parvalbumin-positive (PV^+^) neurons were identified as the most vulnerable in prion diseases [46–49]. In both humans affected by CJDs and mice infected with mouse-adapted strains, a decline in PV^+^ neurons was observed at the subclinical stage of the disease [46–49]. If microglia are responsible for the loss by selectively targeting PV^+^ neurons, we would expect PV^+^ neurons to be engulfed early, i.e., prior to clinical onset. Contrary to this hypothesis, engulfment of PV^+^ neurons was observed only at clinical and terminal time points (Fig. S3A, B). Moreover, both PV^+^ and PV^-^ neurons were subject to engulfment, arguing that this process is not selective toward PV^+^ neurons (Fig. S3A).

### Neurons under engulfment lack apoptotic markers

Our findings suggest that individual engulfment events are arrested or protracted, while partially engulfed neurons remain viable. To investigate this further, we assessed the presence of apoptotic markers in partially engulfed neurons. Staining terminal SSLOW-infected C57Bl/6J mice with an antibody to activated (cleaved) caspase 3 (cCasp3), an early apoptosis marker, revealed that the majority of neurons undergoing engulfment were cCasp3 negative (Fig. 5A,D,E). In fact, the number of cCasp3^+^ neurons was low and comparable to that of normally aged mice (Fig. 5E). Contrary, in positive controls, C57Bl/6J mice subjected to ischemia showed extensive neuronal cCasp3 staining (Fig. 5C). Interestingly, in prion-affected brains, reactive microglia exhibited cCasp3-positive puncta (Fig. 5A,B,E). Confocal microscopy imaging confirmed that cCasp3 immunoreactivity was associated with microglia and displayed intracellular localization (Fig. 5B). Caspase 3 activation in microglia has been recognized as a switch between pro-inflammatory activation and cell death, as observed in neurodegenerative diseases including Alzheimer’s and Parkinson’s diseases [50, 51].

**Figure 5.**
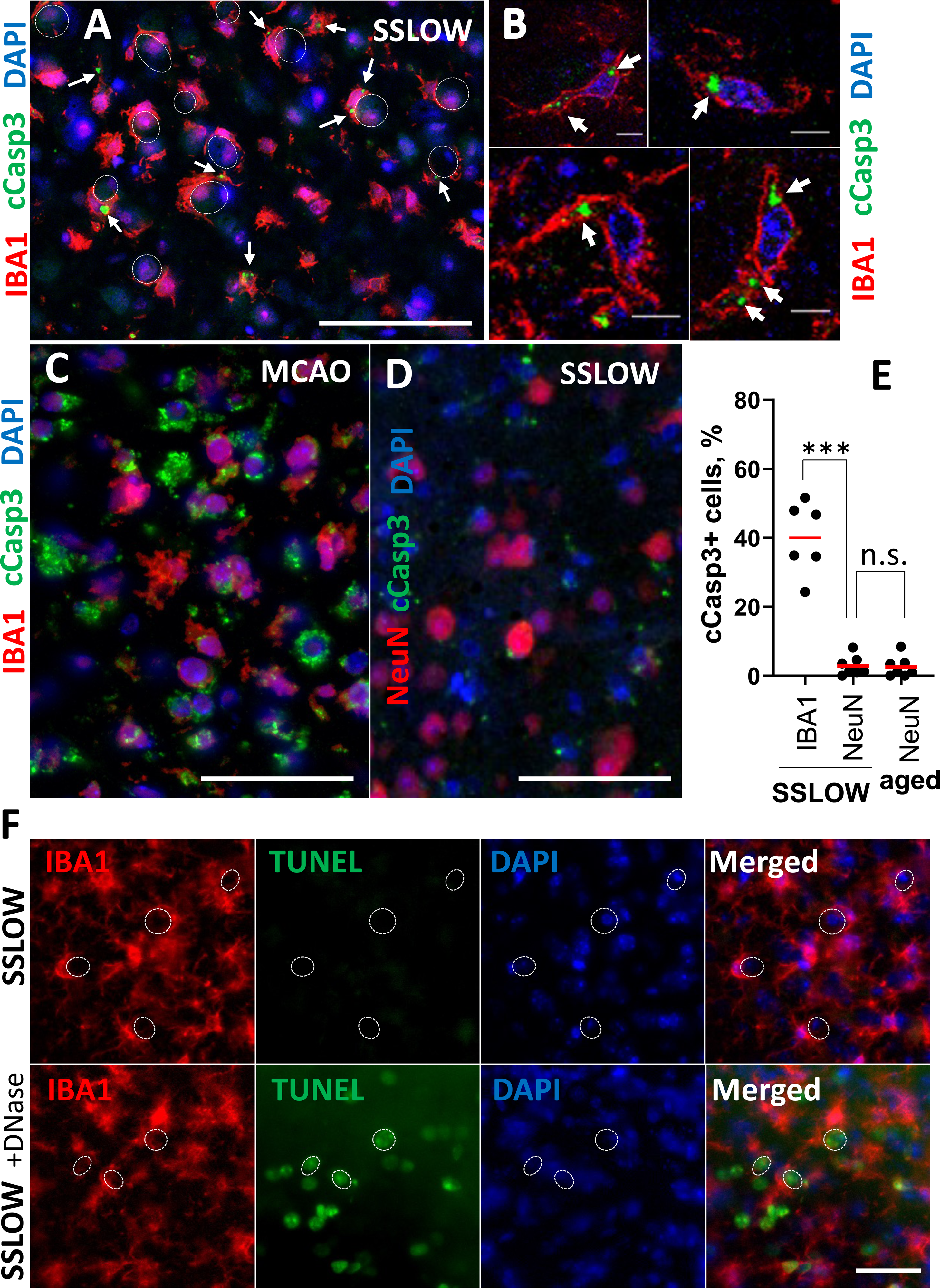
Engulfed neurons lack apoptotic markers. (**A-C**) Coimmunostaining of brain sections of C57Bl/6J mice infected with SSLOW and analyzed at the terminal stage of the disease (**A, B**) or C57Bl/6J mice subjected to MCAO and analyzed 5 days post-insult (**C**) using antibody to activated (cleaved) caspase-3 (cCasp3) and anti-IBA1 antibody. (**B**) Confocal microscopy imaging illustrates the intracellular localization of cCasp3 in microglia of SSLOW-infected mice. Arrows point at cCasp3 puncta. Dashed circular lines represent neuronal engulfment. (**D**) Coimmunostaining of brain sections of C57Bl/6J mice infected with SSLOW using anti-cCasp3 and anti-NeuN antibodies. (**E**) Percentage of cCasp3^+^ microglia (IBA1) and neurons (NeuN) in cortexes of SSLOW-infected mice and in neurons of aged 607-740-day-old C57Bl/6J mice. ***p < 0.001 by unpaired t-test with Welch’s correction. N=6-7 fields of view. (**F**) TUNEL staining of brain sections of a C57Bl/6J mouse infected with SSLOW via i.c. route, and sections from the same mouse pre-treated with DNase and used as positive controls. Dashed circular lines represent examples of neuronal engulfment. Scale bars = 50 µm in A, C and D, 5 µm in B, and 100 µm in F.

As an alternative approach for identifying apoptotic cells in prion-infected C57Bl/6J mice, we employed the terminal transferase dUTP nick end labeling (TUNEL) assay. TUNEL stains apoptotic cells by detecting DNA fragmentation. In the cortices of terminal SSLOW-infected mice, neurons undergoing engulfment were negative for TUNEL staining (Fig. 5F). However, pretreatment of brain slices from the same mice with DNase revealed DNA fragmentation detected by the TUNEL staining. In summary, these results indicate that neurons undergoing engulfment by microglia lack apoptotic markers.

### Neurons are engulfed by PrP^Sc^-positive microglia

Reactive microglia may target viable neurons because they sense PrP^Sc^ on neuronal surfaces. To investigate whether neurons undergoing engulfment are PrP^Sc^-positive, brains of SSLOW-infected C57Bl/6J mice were coimmunostained using anti-PrP antibody 3D17 in combination with anti-MAP2, anti-IBA1, or anti-GFAP antibodies. In SSLOW-infected animals, the majority of PrP immunoreactivity was associated with reactive microglia, including the microglia that engulfed neurons (Fig. 6A,C). Non-infected, age-matched controls displayed very weak PrP immunoreactivity, arguing that in prion-infected animals, the majority of PrP signal is likely to be attributed to PrP^Sc^ (Fig. 6B,F). In SSLOW-infected mice, only a subtle PrP signal was detected in neurons (Fig. 6A,D). Very minimal, if any, PrP immunoreactivity was detected in astrocytes (Fig. 6A,E). A similar pattern of PrP immunoreactivity, the majority of which colocalized with microglia, was observed in а parallel staining using different anti-PrP antibody SAF-84 (Fig. S4). Unlike neurons or astrocytes, microglia do not replicate PrP^Sc^ but can acquire PrP^Sc^-positivity via phagocytic uptake [26, 27].

**Figure 6.**
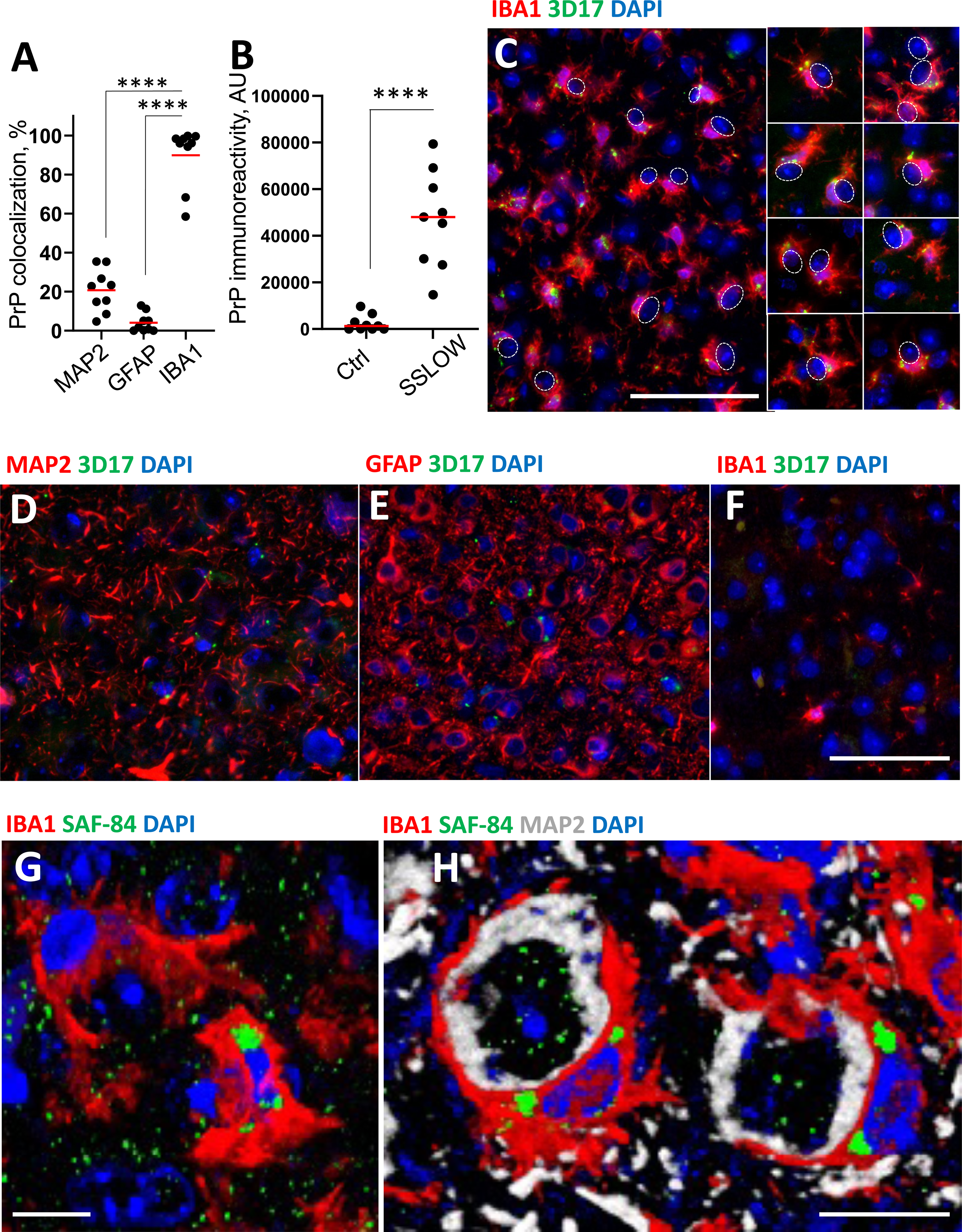
PrP^Sc^ colocalizes with reactive microglia. (**A**) Quantification of PrP^Sc^ colocalization with MAP2^+^, GFAP^+^ and Iba1^+^ cells in C57Bl/6J mice infected with SSLOW and analyzed at the terminal stage of the disease. **(B)** Quantification of PrP immunoreactivity in SSLOW-infected and age-matched control C57Bl/6J mice. In **A** and **B**, N=3 animals per group, n=9 fields of view; **** p<0.0001 by non-parametric Mann-Whitney test. (**C-E**) Representative images of SSLOW-infected C57Bl/6J mouse brains coimmunostained using anti-PrP (3D17) and anti-IBA1 antibodies (**C**), anti-MAP2 (**D**), or anti-GFAP antibodies (**E**) at the terminal stage of the disease. In **C**, dashed circles show neuronal engulfment; a gallery of additional images on the right shows PrP^Sc+^ microglia that engulf neurons. (**F**) Co-immunostaining of non-infected, age-matched C57Bl/6J mouse brains using anti-PrP (3D17) and anti-IBA1 antibodies. (**G, H**) 3D reconstruction of confocal microscopy images of microglia engulfing neurons. Coimmunostaining of SSLOW-infected C57Bl/6J mouse brains at terminal stages of the disease using anti-PrP (SAF-84) and anti-Iba1 antibodies (**G**) or anti-PrP (SAF-84) and anti-Iba1 and anti-MAP2 antibodies (**H**). Scale bars = 50 µm for C – F, and 5 µm in G and H.

Three-dimensional reconstruction of confocal microscopy imaging confirmed the intracellular localization of PrP^Sc^ aggregates in reactive microglia (Fig. 6G). Furthermore, triple coimmunostaining using anti-MAP2 and anti-IBA1 antibodies revealed that PrP^Sc^ aggregates in microglia engulfing neurons localized to perinuclear sites (Fig. 6H and S5). Moreover, confocal microscopy Z-stack video demonstrate deposition of PrP^Sc^ that occupies substantial volume in microglia engulfing neurons (Fig. S5).

### Phagocytic uptake of PrP^Sc^ by microglia precedes neuronal engulfment

The observation of PrP^Sc^ deposits in reactive microglia engaged in neuronal engulfment raises the possibility that microglia phagocytically uptake PrP^Sc^ prior to engulfment. To test this hypothesis, we quantified the percentage of PrP^Sc+^ microglial cells and the percentage of neurons undergoing engulfment in the cortices of C57Bl/6J mice at regular time points starting at 64 dpi. A significant increase in the percentage of PrP^Sc+^ microglia was observed at 78 dpi (Fig. 7A, *i* and S6), whereas engulfment became noticeable only from 106 dpi onward (Fig. 7A, *ii*). At 106 dpi, almost 50% of microglial cells were PrP^Sc+^(Fig. 7A, *i*). Remarkably, the total amount of PrP^Sc^, as quantified by Western blot, remained at very low levels until 106 dpi (Fig. 7A, *iv*), suggesting that microglia manage to control prion replication through their phagocytic uptake. Notably, significant proliferation of microglia was observed after 106 dpi (Fig. 7A, *iii*), tripling their phagocytic capacity. Consistent with a substantial boost in phagocytic capacity was an increase in CD11b and Gal3 levels observed during the clinical stage (Fig. 7 B,C). CD11b and Gal3 are involved in two different phagocytic pathways. Notably, the percentage of PrP^Sc+^ microglia continued to rise even after 106 dpi (Fig. 7A, *i*) alongside their substantial proliferation (Fig. 7A, iii), indicating that proliferated cells were actively engaged in the phagocytic uptake of PrP^Sc^ (Fig. 7A, *v*). However, despite the significant boost in phagocytic capacity attributed to proliferation, prion replication spun out of control after 106 dpi (Fig. 7A, *iv* and B). This time point coincides with the initiation of а microglial encircling of neurons, which only intensifies with the disease progression (Fig. 7A, *ii*). To summarize, phagocytic uptake of PrP^Sc^ by microglia preceded neuronal engulfment by four weeks, whereas clinical onset followed neuronal engulfment. These findings suggest that PrP^Sc^ uptake upregulates phagocytic activity and, perhaps, primes microglia for initiating the engulfment of neurons.

**Figure 7.**
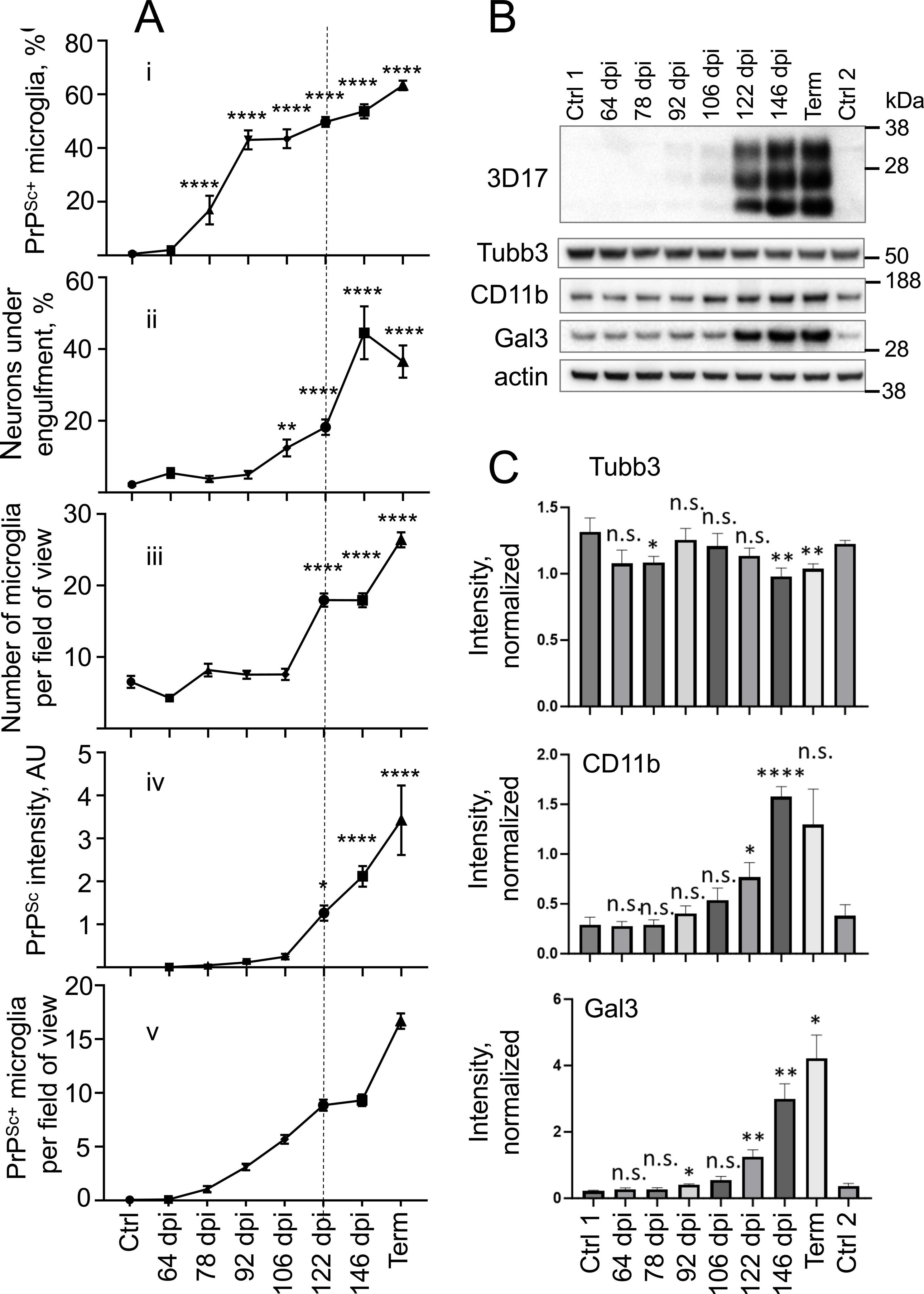
Time course of PrP^Sc^ uptake, neuronal engulfment, and microglia proliferation. (**A**) Changes in the percentage of PrP^Sc+^ microglia (***i***), the percentage of MAP^+^ neurons under engulfment (***ii***), the total number of IBA1^+^ microglial cells per field of view (***iii***), the amounts of PrP^Sc^ as estimated by Western blot (**iv**), and the total number of PrP^Sc+^ microglia per field of view (***v***) in C57Bl/6J mice infected with SSLOW via i.p. route during subclinical and clinical stages of the disease. For each time point, N=3-5 individual animals, n=18-32 fields of view for plots shown in ***ii*,** and N=4-5 individual animals, n**=**17-60 for ***i, iii*** and ***v***. * p<0.05, ** p<0.01, **** p<0.0001 by unpaired Mann-Whitney’s test. Ctr: age-matched control for 64 dpi and terminal points. Term: terminal animals collected at 157-166 dpi. **(B)** Representative Western blots of PrP^Sc^, Tubb3, CD11b and Gal3 in brain material of SSLOW-infected C57Bl/6J mice during preclinical and clinical stages of the diseases. (**C**) Quantification of Western blots of Tubb3, CD11b and Gal3. Signal intensities of Tubb, CD11b and Gal3 were normalized per intensities of actin for each individual Western blot. In **B** and **C**, Ctrl1: age-matched control for 64 dpi. Ctrl2: age-matched control for terminal time point. Term: terminal animals collected at 157-166 dpi. N=3-5 animals. Data presented as mean ± SEM. * p<0.05, ** p<0.01, **** p<0.0001; each time point was compared to the combined control group (Ctrl1 + Ctrl2) by an unpaired t-test with Welch’s corrections.

### Microglia engaged in engulfment are characterized by hypertrophic lysosomal compartments

The sustained uptake of PrP^Sc^ during the preclinical stage appears to prime microglia for their engulfment of neurons. Given that sustained phagocytic activity necessitates the upregulation of lysosomal degradation, we proceeded to quantify lysosomal compartments in individual microglial cells through the confocal microscopy imaging of Lysosome Associated Membrane Protein 1 (LAMP1). LAMP1, which is highly expressed in lysosomal membranes, served as a lysosomal marker. In contrast to age-matched controls, in SSLOW-infected animals, we observed a significant LAMP1 immunoreactivity associated with microglia (Fig. 8A). Remarkably, within the SSLOW-infected brains, LAMP1 immunoreactivity was notably higher in microglial cells engaged in neuronal engulfment compared to microglia not involved in engulfment (Fig. 8B, C). These results support the idea that microglial cells attempting to engulf neurons are primed for lysosomal degradation.

**Figure 8.**
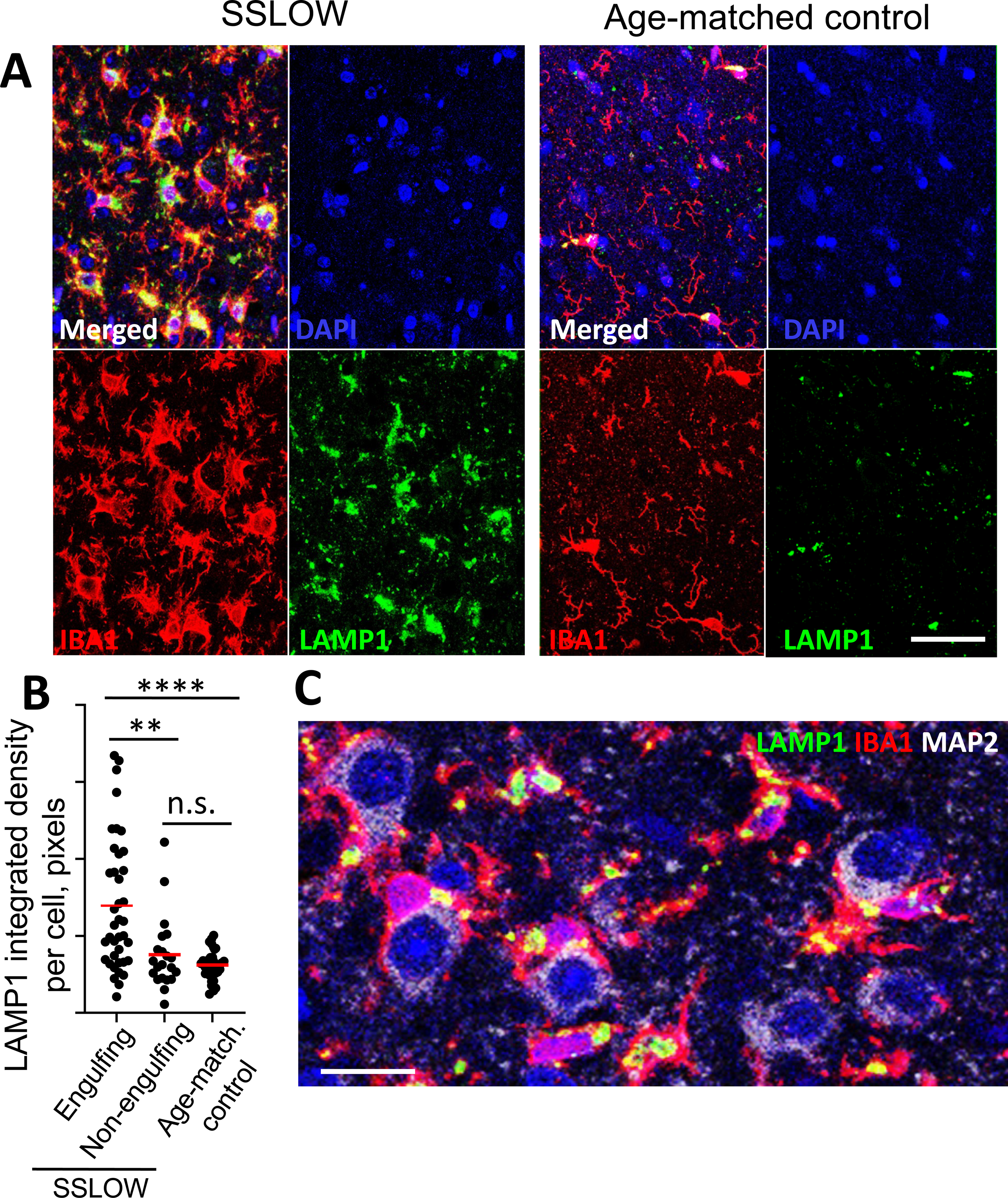
Hypertrophic lysosomes in microglia engaged in engulfment. (**A**) Representative maximum intensity projection confocal images of LAMP1^+^ compartments in cortices of C57Bl/6J mice infected with SSLOW and analyzed at the terminal stage of the disease along with age-matched control mice. Brain sections were coimmunostained using anti-LAMP1 (green) and anti-IBA1(red) antibodies. (**B**) Quantification of LAMP1 integrated density in individual microglial cells engaged or not engaged in neuronal engulfment in cortices of SSLOW-infected mice and age-matched controls. Integrated density was analyzed using maximum intensity projection 3D images from confocal microscopy. n=21-39 individual cells, ** p<0.01, *** p<0.001 by unpaired Mann-Whitney’s test. (**C**) Representative confocal microscopy image of LAMP1^+^ compartments in cortices of C57Bl/6J mice infected with SSLOW. Scale bars = 25 µm in A, and 10 µm in C.

### Neuronal engulfment is independent of CD11b pathway

Among several microglial phagocytic pathways, CD11b-dependent pathway was previously found to be responsible for phagocytosis of newborn cells, including neurons during development [42, 52]. CD11b was also shown to be involved in the phagocytosis of neurons in glia-neuronal co-cultures [53]. To test whether the same pathway is responsible for engulfment in prion-infected mice, we analyzed the time-course of the disease and neuronal engulfment in CD11b knock-out mice infected with SSLOW strain (Fig. 9). There were no differences with respect to incubation time to the terminal disease or the amount of PrP^Sc^ between CD11b^−/−^ and control C57Bl/6J (WT) groups (Fig. 9A-C). Neurons were under engulfment in both, CD11b^−/−^ and control groups (Fig. 9D), while the percentage of engulfed neurons was the same in the two groups (Fig. 8E). Moreover, no differences between CD11b^−/−^ and control WT group were found with respect to the density of MAP2^+^ neurons (Fig. 9E), the level of expression of neuron-specific protein Tubb3 (Fig. 9B,C), or microglia activation, as judged from IBA1 immunoreactivity (Fig. 9E). Modest upregulation of galectin 3 (Gal3) was seen in CD11b^−/−^ versus control group (Fig. 9B,C). Gal3, which is involved in alternative phagocytic pathways, is released by activated myeloid cells and acts as an opsonin by binding to galactose residues on the cell surface.

**Figure 9.**
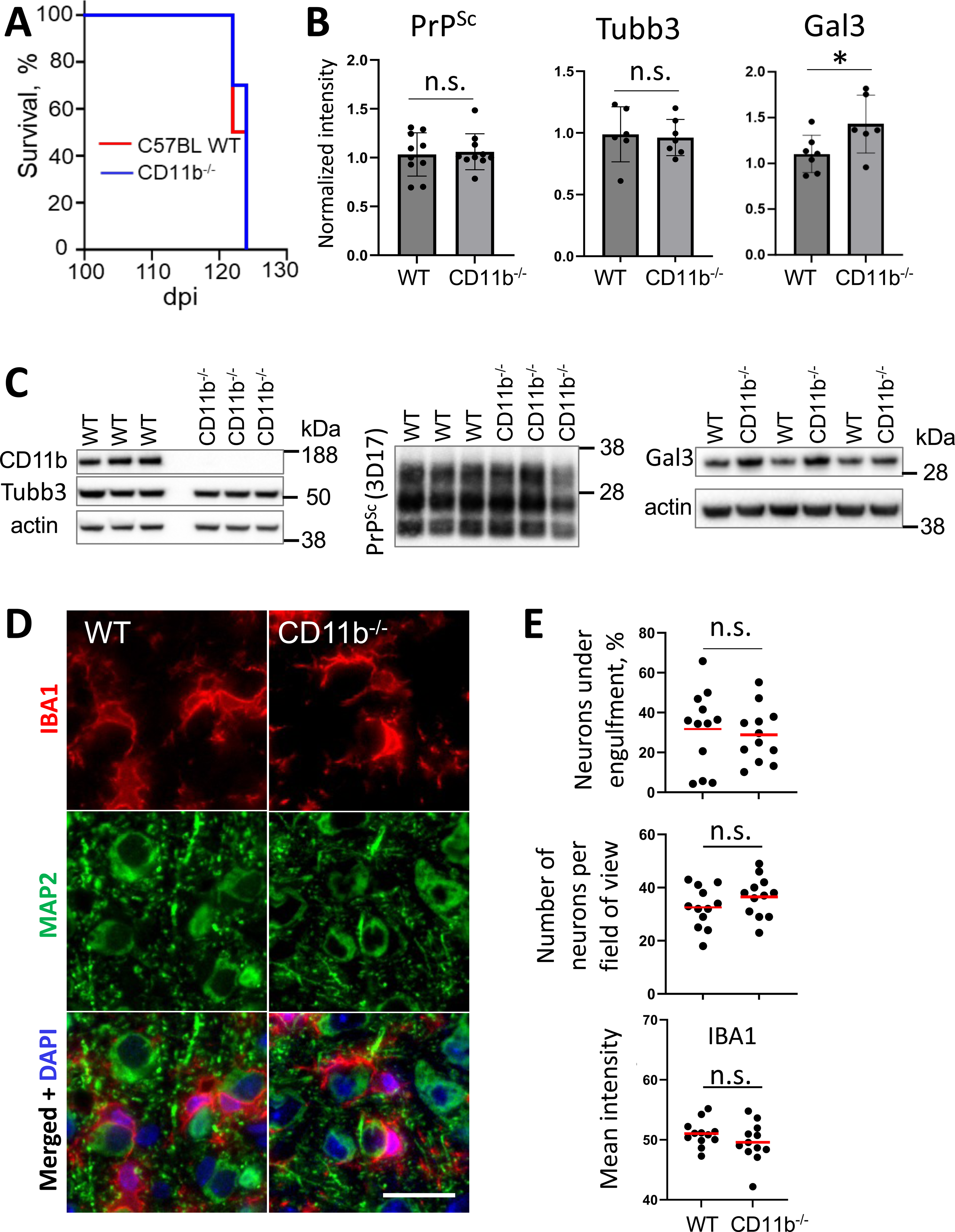
Analysis of prion disease pathogenesis and neuronal engulfment in CD11b^−/−^ mice. (**A**) Incubation time to terminal disease in CD11b^−/−^ and C57Bl/6J control mice (WT) inoculated with SSLOW via i.c. route. N=10 individual animals for each group. Mantel-Cox test of survival curves indicated no significant difference between CD11b−/− and WT groups. (B) Densitometric quantification of Western blots for PrP^Sc^, Tubb3 and Gal3 in CD11b^−/−^ and WT control mice infected with SSLOW. Data represent means ± SD, N=10 for each group, * *p*<0.05, and ‘n.s.’ non-significant by two-tailed, unpaired Student’s t-test. (**C**) Representative Western blots of selected markers for brains of CD11b^−/−^ and C57Bl/6J control mice (WT) at the terminal stage of the diseases. For analysis of PrP^Sc^, BHs were digested with PK, and blots were stained with 3D17 antibody. (**D**) Representative images of partial engulfment of neurons by microglia in the cortex of CD11b^−/−^ and WT control mice infected with SSLOW and analyzed at the terminal stage of the disease. Brain sections were stained using anti-IBA1 (red) and anti-MAP2 (green) antibodies. (**E**) Percentage of MAP2^+^ neurons undergoing engulfment, the total number of MAP2^+^ neurons, and quantification of IBA1 immunoreactivity in CD11b^−/−^ and WT control mice infected with SSLOW at terminal stages of the disease. N=4 individual animals per group, n=12 fields of view. Means are marked by solid red lines. n.s. – no significant differences by unpaired Student’s t-test. Scale bar = 20 µm.

### Neuronal engulfment in sCJD individuals

To check whether the partial engulfment phenomenon occurs in human prion diseases, we examined sCJD brains using staining for β-chain of human HLA-DR, a marker of reactive microglia. In all examined sCJD subtypes, including MM1, MM2C, VV1, VV2 and MV2K [54], we found reactive microglia engaged in partial engulfment (Fig. 10). Akin to prion-infected mouse brains (Fig. 10A), such partial engulfment was detected across different human brain regions affected by prions, including frontal and temporal cortex, thalamus, striatum, midbrain and hippocampus (Fig. 10B-F).

**Figure 10.**
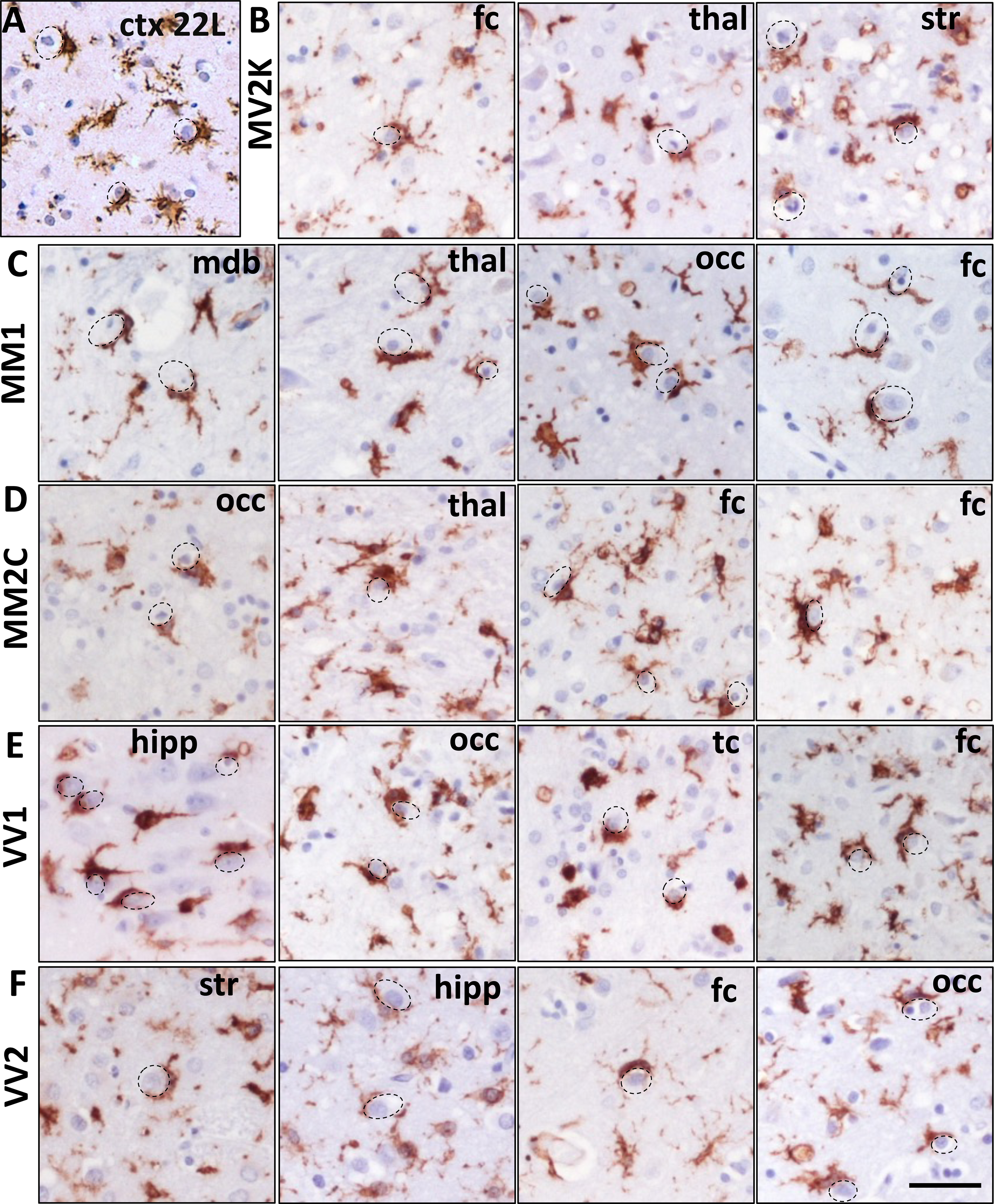
Partial engulfment by reactive microglia in sCJD. (**A**) Representative image of reactive microglia stained with anti-IBA1 antibody in the cortex (ctx) of 22L-infected C57Bl/6J mice provided as a reference. **(B-F)** Representative images of reactive microglia strained with anti-HLA-DR+DP+DQ [CR3/43] antibody in the following subtypes of sCJD: MV2K (**B**), MM1 (**C**), MM2C (**D**), VV1 (**E**) and VV2 (**F**). Fc, frontal cortex; thal, thalamus; str, striatum; mdb, midbrain; occ, occipital cortex; hipp, hippocampus; tc, temporal cortex. Dashed circular lines indicate neuronal engulfment. Scale bar = 20 µm.

## Discussion

The present study describes a novel phenomenon of partial engulfment of neurons by reactive microglia. Partial engulfment was consistently observed across mouse-adapted strains in mice and in different subtypes of sCJD in humans. Furthermore, this partial engulfment phenomenon manifested in multiple brain regions affected by prions.

Several lines of evidence suggest that the engulfment process is either stalled or aborted. First, a mere fraction of less than one percent of neurons were found to be fully engulfed. Second, although a significant proportion of neurons exhibited engulfment during the clinical stage of the disease (Fig. 4B), this process was not accompanied by neuronal loss. Third, considering that complete phagocytosis of a cell in a mouse brain takes only 25 minutes to 2 hours [15, 45], the entire neuronal population should theoretically have been phagocytosed within less than 24 hours, assuming 10% of neurons undergoing engulfment at any given time point. However, in prion-infected mice, engulfment persisted at the rate of ten percent or above for several weeks, while neuronal density did not decrease. These findings collectively indicate a unique pattern of incomplete engulfment in the context of neurodegenerative diseases and challenge conventional expectations of the temporal dynamics of microglial phagocytosis.

It is unclear whether engulfment was irreversibly aborted, locked in a dormant stage (i.e. potentially reversible), or simply protracted. Nevertheless, extensive sequestration of neuronal surfaces by microglia is likely to induce neuronal dysfunction even without neuronal death. This is evident from a significant downregulation of neuron-specific genes responsible for synaptic neurotransmission, memory, and learning. Additionally, it remains unclear whether the failure of phagocytosis of neurons can be attributed to a specific state of reactive microglia that cannot complete the engulfment process. The premature termination of phagocytosis could have resulted from overloading microglia with PrP^Sc^, driving microglia to a senescent state and affecting their main function of phagocytosis. Supporting this hypothesis, previous studies have shown that microglia acquire a senescent phenotype in cultures upon phagocytosing live neurons with tau aggregates [11, 12]. Notably, the observation of cCasp3 in reactive microglia in the current work is consistent with activation-induced apoptosis described as one of the consequences of microglial activation [55]. In fact, in Alzheimer’s and Parkinson’s diseases, activation of caspase 3 in microglia was found to regulate the switch between pro-inflammatory activation and cell death [50, 51]. An alternative hypothesis regarding incomplete engulfment proposes that engulfment serves a neuroprotective role and is meant to be partial. If this is the case, microglia-to-neuron contacts could be dynamic. One could hypothesize that activated microglia survey neurons via encircling neuronal surfaces and establishing temporal contacts followed by their decoupling. However, it is challenging to reconcile this hypothesis with the fact that microglial cells that engaged in engulfment are primed for phagocytosis, as evident from hypertrophic lysosomes and upregulated phagocytic pathways [56]. Alternatively, a cross-talk between microglia and neurons [57], which appear to be still viable while partially engulfed, might contribute to engulfment arrest. Neurons might employ an unknown mechanism that resists microglial attempts to phagocytose them. These findings underscore the significance of understanding the intricate interplay between microglial phagocytosis and neuronal involvement in the pathogenesis of prion diseases.

The present study provides new important insight into the timeline of disease pathogenesis. The enhanced stability of SSLOW PrP^Sc^ within the phagocytic compartments of microglia, compared to PrP^Sc^ from other strains, provides unique opportunities for quantitatively assessing PrP^Sc^ phagocytosis in the brain [58]. During the preclinical stage, microglia appear to keep PrP^Sc^ replication at low leveel. Notably, the percentage of PrP^Sc+^ microglial cells increases rapidly during the preclinical stage (Fig. 7A, *i*), when the total amount of PrP^Sc^ still remains at low levels (Fig. 7A, *iv* and B). This dynamic suggests that the potential accumulation of PrP^Sc^ resulting from replication is counteracted by the phagocytosis and clearance of PrP^Sc^. The proliferation of microglia, particularly evident after 106 dpi, triples their phagocytic capacity (Fig. 7A, *iii*). Notably, the percentage of PrP^Sc+^ microglia continues to rise even after 106 dpi (Fig. 7A, *i*) in conjunction with their active proliferation (Fig. 7A, *iii*), suggesting that proliferated cells are competent in the phagocytic uptake of PrP^Sc^. However, despite the considerable increase in the total phagocytic capacity (Fig. 7A, *v*), the rate of PrP^Sc^ replication surpasses the rate of its clearance, leading to a steady buildup of PrP^Sc^ after 106 dpi (Fig. 7A, *iv* and B).

Remarkably, the late preclinical stage, spanning from 92 to 106 dpi, marks a critical time window when the phagocytic uptake of PrP^Sc^ slows down. Instead, microglia shift their focus and engage in engulfment of neurons (Fig. 7A*i*, *ii*). It remains unclear whether this change in microglial behavior is simply attributed to the hyperactivation of the phagocytic machinery in general and/or dysregulation in competitive pathways responsible for the phagocytosis of different targets. Nevertheless, our studies reveal that PrP^Sc^ uptake by microglia precedes their engulfment of neurons, whereas clinical onset follows neuronal engulfment. Based on this dynamic, we proposed that phagocytosis of PrP^Sc^ primes microglia for engulfment of live neurons by upregulating phagocytic pathways (Fig. 11). Supporting this mechanism, microglial cells engaged in neuronal engulfment were found to be PrP^Sc^-positive and exhibited hypertrophic lysosomes, indicating their priming for phagocytosis. Consistent with the hypothesis mentioned above are recent studies demonstrating that microglia are activated directly by PrP^Sc^ [59]. Moreover, phagocytic activity was significantly upregulated in microglia isolated from prion-infected animals [56].

**Figure 11.**
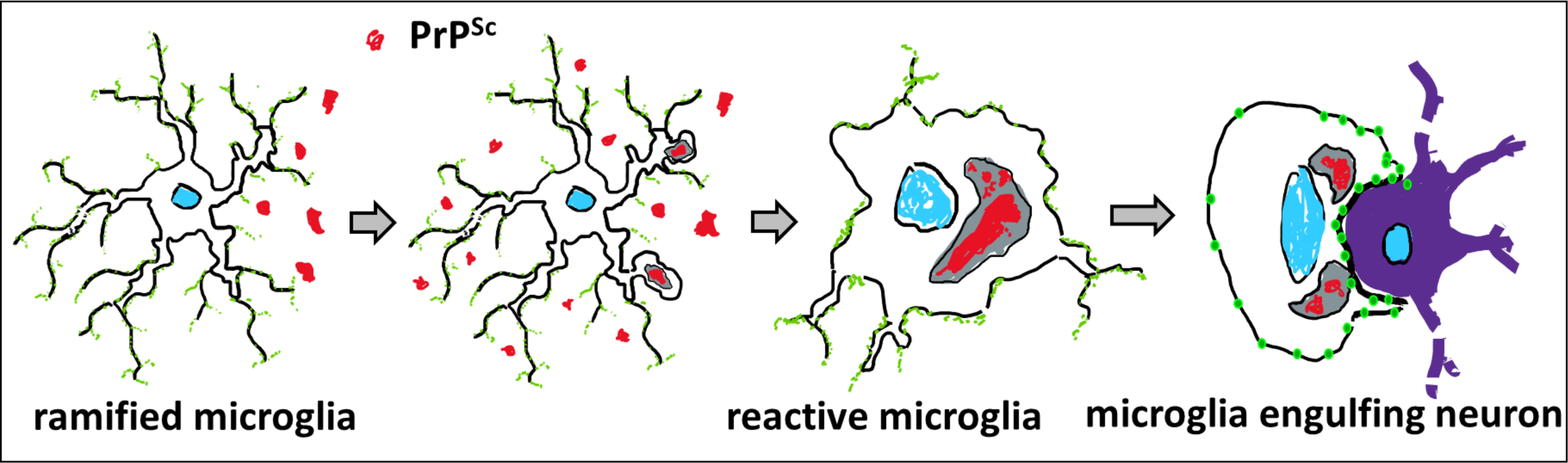
Schematic diagram illustrating that uptake of PrP^Sc^ by microglia precedes engulfment of neurons. Phagocytic uptake of PrP^Sc^ activates phagocytic pathways and primes microglia for their attack on neurons.

Do neurons exhibit any cues that provoke a phagocytic attack by microglia? The hypothesis that reactive microglia selectively engulf GABAergic, PV^+^ neurons – previously identified as the most vulnerable to prion infection [46–49] – was not supported by our experimental results. The subpopulation of PV^+^ neurons was not among the earliest targets by microglia. In the later stages of the disease, they were under engulfment along with PV^-^ neurons. As an alternative hypothesis, microglia could target neurons because the phagocytes sense PrP^Sc^ on neuronal surfaces. In mice infected with SSLOW, the majority of PrP^Sc^ was detected in microglia, yet PrP immunoreactivity was also observed in association with neurons. Unlike microglia, neurons do express PrP^C^. Because attributing neuronal PrP immunoreactivity in individual cells exclusively to PrP^Sc^ or PrP^C^ is challenging, it is difficult to prove or dismiss the hypothesis of neuronal PrP^Sc^ as a cue for microglia. Nevertheless, the fact that neuronal engulfment is shared between different mouse-adapted strains (Fig. 2), regardless of their cell tropism, suggests that the presence of PrP^Sc^ on the neuronal surface might not be the major factor driving engulfment. In fact, in mice infected with ME7, PrP^Sc^ is reported to be predominantly associated with neurons, whereas in 22L-infected mice, the majority of PrP^Sc^ is associated with astrocytes [44]. Yet the percentage of neuronal engulfment was the same in these two strains. The third hypothesis proposes that neurons enter apoptotic cell death, and apoptosis-associated cues activate an eat-me signaling in microglia, resulting in engulfment. However, the absence of activated caspase 3, a marker of early apoptosis, and TUNEL staining in neurons undergoing engulfment suggests that they were not apoptotic. Moreover, neuronal density did not decline (Fig. 4A), despite a higher than twenty percent engulfment during the clinical stage that lasted one month (Fig. 7A). These findings argue that partially engulfed neurons remain viable. The lack of neuronal loss is in agreement with previous reports by Aguzzi and coauthors, who attributed the lack of neuronal death to the relatively early endpoint imposed by animal welfare regulations [60]. 20% weight loss, used here as an endpoint, happens when animal’s motor impairment affects its ability to reach food and water in a cage. We do not know how long mice would live if artificially supported or after a 20% weight loss and whether they would experience any neuronal loss upon approaching the true endpoint.

Together with CD18, CD11b constitutes complement receptor 3 (CR3). The CR3-dependent phagocytic pathway involves the complement factor C3b, which tags neurons and synapses, driving phagocytosis through interaction with CR3 [2, 3, 8]. The CD11b-dependent pathway has been identified as responsible for the phagocytosis of neurons during development [42, 52]. Furthermore, CR3-dependent mechanisms have been implicated in the elimination of synapses and neurons in Alzheimer’s disease, frontotemporal dementia, and normal aging [8, 9, 61, 62]. A significant upregulation of *C3* and *Itgam*, the gene encoding CD11b, was observed in prion-infected mice and sCJD individuals [58, 63]. However, the knockout of CD11b in the current study did not affect the engulfment of neurons by microglia, nor did it alter the incubation times to terminal disease. These findings suggest that a CD11b-independent pathway orchestrates the incomplete engulfment of neurons in prion diseases.

An upregulation of Gal3 with disease progression (Fig. 7B,C), along with an additional upregulation upon CD11b knockout, points to the Gal3-dependent pathway as a plausible mechanism responsible for neuronal engulfment in prion diseases (Fig. 9C,D). Gal3, released by activated myeloid cells, acts as an opsonin by binding to galactose residues of N-linked glycans lacking terminal sialic acid residues (43,44). Several lines of evidence suggest that galactose on PrP^Sc^ N-linked glycans is responsible for microglia activation. Partial desialylation of PrP^Sc^, resulting in an increase in the amounts of exposed galactose, has been found to enhance the inflammatory response of microglia in cell cultures [59]. Prion strains recruit PrP^C^ sialoglycoforms selectively, according to strain-specific structure [64, 65]. Consequently, prion strains are characterized by different ratios of galactose versus sialic acid residues on the PrP^Sc^ surface [66]. Among four mouse-adapted strains employed here, SSLOW exhibits the highest level of exposed galactose, causing the most profound and widespread neuroinflammation, and has the shortest disease incubation time [58]. In contrast, ME7 PrP^Sc^ displays the highest sialylation level, presenting the most attenuated neuroinflammation and the longest incubation time [58]. In addition to the strain-specified differences, PrP^Sc^ sialylation is also controlled by brain region [67]. PrP^Sc^ produced in the thalamus is less sialylated than PrP^Sc^ from the hippocampus or cortex [67]. Remarkably, the thalamus is the first to develop neuroinflammation and is the most severely affected at the terminal stage [41, 67–69]. Based on these findings, we propose that PrP^Sc^ activates phagocytic pathways in microglia that sense exposed galactose as an ‘eat-me’ signal.

It has been shown that during brain development, microglia can phagocytose viable non-apoptotic cells, including neural precursor cells [45, 70]. Moreover, microglial attacks on viable non-apoptotic cells have also been suggested in neurodegenerative diseases (reviewed in [7]). In inherited retinal degeneration, microglia have been shown to phagocytose non-apoptotic photoreceptor cells [71]. Surprisingly, injections of Aβ into the mouse brain were found to induce the phagocytosis of neurons by microglia, suggesting that one-time stimulation might be sufficient to trigger microglial phagocytic activity toward neurons [72]. The present study is the first to illustrate microglial attacks on non-apoptotic neurons in prion diseases. To our knowledge, this study is the first to document the phenomenon of incomplete or stalled engulfment of neurons.

The attack on viable neurons presented here reinstates the debate about the protective versus neurotoxic role of microglia. The uptake of PrP^Sc^ by microglia during the early stages of the disease, as shown here, supports the hypothesis of the protective role of microglia. Previously, ablation of microglia either before prion infection or during the early stages of the disease was found to significantly accelerate disease progression, establishing their protective role [29–32]. However, partial inhibition of microglia proliferation and reactivity at the late preclinical stage delayed disease onset and extended survival by 26 days [34]. Moreover, therapeutic inhibition of microglia activation just before or after the disease onset prolonged the survival of humanized mice infected with sCJD [35]. The present study points to the transformation of the microglial role from predominantly positive during the early stage to a net negative during the late stages of the disease. We propose that this pivotal shift occurs at the late preclinical stage, where neuronal engulfment surpasses the phagocytic uptake of PrP^Sc^. Sustained phagocytic activity, initially a defensive response to prion infection, may eventually become detrimental due to the upregulation of phagocytic pathways that lead to the assault on viable neurons. Consistent with this view, our recent studies documented significant upregulation of phagocytic activity in reactive microglia associated with prion diseases [56]. Moreover, phagocytic microglia did not discriminate between synaptosomes purified from prion-infected brains and normal synaptosomes as phagocytic substrates. Our current work suggests that the timing of the shift in phagocytic activity is crucial, emphasizing the need for a comprehensive understanding of the intricate interplay between microglial phagocytosis and the progression of prion diseases. Importantly, the determination of whether the selectivity of microglial phagocytosis can be regulated to target only specific phagocytic substrates represents a promising avenue for future studies.

## Acknowledgemnts

We thank Dr. Alexey Shevelkin for discussion of experimental procedure of confocal imaging.

## Supplementary Figure Legends

**Figure S1. Engulfing microglia in prion-infected C57Bl/6J mice and 5XFAD mice. (A, B)** Reactive microglia in mice infected with SSLOW via i.p. route at the terminal stage of the disease engulf neuronal soma. Staining was performed using anti-IBA1 (red) and anti-MAP2 (green) antibodies. Dashed circles mark nuclei. Arrows and arrowheads point at neuronal and microglial nuclei, respectively. (**C**) In 10-month-old 5XFADs, microglia form pseudopodia with phagocytic cups that fully engulf neurons. Staining performed using anti-IBA1 (green) and anti-CD68 (red) antibodies. The arrow points at the phagocytic cup. Scale bars = 20 µm.

**Figure S2. Downregulation of genes responsible for neuronal function in prion disease.** Normalized expression of genes responsible for neuronal function in C57Bl/6J mice infected with SSLOW via i.p. route or normal age-matched control animals analyzed by NanoString. Data are presented as box-and-whisker plots, where the midline denotes the median, the x represents the mean, and the ends of the box plot denote the 25th and 75th percentiles. N=9 for controls, N=6 for SSLOW-infected mice. **** p < 0.0001; *** p < 0.001; ** p < 0.01; * p < 0.05 by unpaired Student’s t-test.

**Figure S3. Engulfment does not selectively target PV^+^ neurons. (A)** Coimmunostaining of cortexes of terminally ill C57Bl/6J mice infected i.p. with SSLOW, using anti-IBA1 and anti-PV antibodies. Arrows indicate examples of partially engulfed PV^+^ neurons; not engulfed PV^+^ neurons are marked by arrowheads. Examples of microglia engulfing PV-negative cells are marked by asterisks. **(B)** The percentage of PV^+^ neurons undergoing engulfment (upper plot) and number of PV^+^ neurons (lower plot) at the preclinical stage (106 dpi), clinical onset (122 dpi) and terminal stage of the disease (157-166 dpi) in the cerebral cortex of mice inoculated i.p. with SSLOW. N=3-4 individual animals for each time point, n=20-29 total fields of view. * p<0.05, ** p<0.01, **** p<0.0001 by non-parametric Mann-Whitney test. Scale bar = 50 µm.

**Figure S4. PrP^Sc^ colocalizes with reactive microglia.** (**A-E**) Coimmunostaining of SSLOW-infected C57Bl/6J mouse brains using anti-PrP (SAF-84) and anti-IBA1 antibodies at the preclinical (**A**) and terminal stages of the disease (**B**), or SAF-84 and anti-MAP2 (**C**) or anti-GFAP antibodies (**D**) at the terminal stage of the disease. Circles show neuronal engulfment. (**E**) Co-immunostaining of non-infected, age-matched C57Bl/6J mouse brains using SAF-84 and anti-IBA1 antibodies. Scale bar = 25 μm.

**Figure S5. Confocal microscopy Z-stack video showing engulfment of neurons by PrP^Sc^-positive reactive microglia.** SSLOW-infected C57Bl/6J mouse brains collected at the terminal stage of the disease and coimmunostained using anti-PrP (SAF-84), anti-IBA1 and anti-MAP2 antibodies. Arrows point at intracellular PrP^Sc^ deposits in microglia engulfing neurons. Scale bar = 10 µm.

**Figure S6. PrP^Sc^ colocalizes with reactive microglia.** Representative images of PrP^Sc^ colocalized with IBA1^+^ cells in C57Bl/6J mice infected with SSLOW via i.p. route at the preclinical (78 dpi), onset of symptoms (122 dpi), and clinical stages of the disease (146 dpi). Brain sections were stained with anti-PrP antibody 3D17 and anti-IBA1 antibody. Scale bar = 50 µm.

## Notes

### Competing Interest Statement

The authors have declared no competing interest.

## References

1. Prater KE, Green KJ, Mamde S, Sun W, Cochoit A, Smith CL, Chiou KL, Heath L, Rose SE, Wiley J et al: Human microglia show unique transcriptional changes in Alzheimer’s disease. Nature Aging 2023, 3(7):894–907.

2. Stevens B, Allen NJ, Zazquez LE, Howell GR, Christopherson KS, Nouri N, Micheva KD, Mehalow AK, Huberman AD, Stafford B et al: The classical complement cascade mediates CNS synapse elimination. Cell 2007, 131(6):1164–1178.

3. Schafer Dorothy P, Lehrman Emily K, Kautzman Amanda G, Koyama R, Mardinly Alan R, Yamasaki R, Ransohoff Richard M, Greenberg Michael E, Barres Ben A, Stevens B: Microglia Sculpt Postnatal Neural Circuits in an Activity and Complement-Dependent Manner. Neuron 2012, 74(4):691–705.

4. Keren-Shaul H, Spinrad A, Weiner A, Matcovitch-Natan O, Dvir-Szternfeld R, Ulland TK, David E, Baruch K, Lara-Astaiso D, Toth B et al: A Unique Microglia Type Associated with Restricting Development of Alzheimer’s Disease. Cell 2017, 169(7):1276–1290.e1217.

5. Krasemann S, Madore C, Cialic R, Baufeld C, Calcagno N, El Fatimy R, Beckers L, O’Loughlin E, Xu Y, Fanek Z et al: The TREM2-APOE Pathway Drives the Transcriptional Phenotype of Dysfunctional Microglia in Neurodegenerative Diseases. Immunity 2017, 47(3):566–581.e569.

6. Grubman A, Choo XY, Chew G, Ouyang JF, Sun G, Croft NP, Rossello FJ, Simmons R, Buckberry S, Landin DV et al: Transcriptional signature in microglia associated with Aβ plaque phagocytosis. Nature Communications 2021, 12(1):3015.

7. Butler CA, Popescu AS, Kitchener EJA, Allendorf DH, Puigdellívol M, Brown GC: Microglial phagocytosis of neurons in neurodegeneration, and its regulation. J Neurochem 2021, 158(3):621–639.

8. Hong S, Beja-Glasser VF, Nfonoyim BM, Frouin A, Li S, Ramakrishnan S, Merry KM, Shi Q, Rosenthal A, Barres BA et al: Complement and microglia mediate early synapse loss in Alzheimer mouse models. Science 2016, 352(6286):712–716.

9. Stephan AH, Madison DV, Mateos JM, Fraser DA, Lovelett EA, Coutellier L, Kim L, Tsai H-H, Huang EJ, Rowitch DH et al: A Dramatic Increase of C1q Protein in the CNS during Normal Aging. The Journal of Neuroscience 2013, 33(33):13460–13474.

10. Fourgeaud L, Través PG, Tufail Y, Leal-Bailey H, Lew ED, Burrola PG, Callaway P, Zagórska A, Rothlin CV, Nimmerjahn A et al: TAM receptors regulate multiple features of microglial physiology. Nature 2016, 532:240.

11. Brelstaff JH, Mason M, Katsinelos T, McEwan WA, Ghetti B, Tolkovsky AM, Spillantini MG: Microglia become hypofunctional and release metalloproteases and tau seeds when phagocytosing live neurons with P301S tau aggregates. Sci Adv 2021, 7(43):eabg4980.

12. Brelstaff J, Tolkovsky AM, Ghetti B, Goedert M, Spillantini MG: Living Neurons with Tau Filaments Aberrantly Expose Phosphatidylserine and Are Phagocytosed by Microglia. Cell Rep 2018, 24(8):1939–1948.e1934.

13. Nomura K, Vilalta A, Allendorf DH, Hornik TC, Brown GC: Activated Microglia Desialylate and Phagocytose Cells via Neuraminidase, Galectin-3, and Mer Tyrosine Kinase. J Immunol 2017, 198(12):4792–4801.

14. Sierra A, Abiega O, Shahraz A, Neumann H: Janus-faced microglia: beneficial and detrimental consequences of microglial phagocytosis. Front Cell Neurosci 2013, 7:6.

15. Sierra A, Encinas JM, Deudero JJ, Chancey JH, Enikolopov G, Overstreet-Wadiche LS, Tsirka SE, Maletic-Savatic M: Microglia shape adult hippocampal neurogenesis through apoptosis-coupled phagocytosis. Cell Stem Cell 2010, 7(4):483–495.

16. Prusiner SB: Prions. Proc Natl Acad Sci U S A 1998, 95:13363–13383.

17. Prusiner SB: Novel proteinaceous infectious particles cause scrapie. Science 1982, 216(4542):136–144.

18. Cohen FE, Prusiner SB: Pathologic conformations of prion proteins. Annu Rev Biochem 1998, 67:793–819.

19. Fang C, Imberdis T, Garza MC, Wille H, Harris DA: A Neuronal Culture System to Detect Prion Synaptotoxicity. PLoS Pathog 2016, 12(5):e1005623.

20. Fang C, Wu B, Le NTT, Imberdis T, Mercer RCC, Harris DA: Prions activate a p38 MAPK synaptotoxic signaling pathway. PLOS Pathogens 2018, 14(9):e1007283.

21. Solforosi L, Criado JR, McGavern DB, Wirz S, Sanchez-Alavez M, Sugama S, DeGiorgio LA, Volpe BT, Wiseman E, Abalos G et al: Cross-Linking Cellular Prion Protein Triggers Neuronal Apoptosis in Vivo. Science 2004, 303:1514–1516.

22. Brandner S, Isenmann S, Raeber A, Fischer M, Sailer A, Kobayashi Y, Marino S, Weissmann C, Aguzzi A: Normal host prion protein necessary for scrapie-induced neurotoxicity. Nature 1996, 379:339–343.

23. Mallucci G, Dickinson A, Linehan J, Klohn PC, Brandner S, Collinge J: Depleting Neuronal PrP in Prion Infection Prevents Disease and Reverses Spongiosis. Science 2003, 302:871–874.

24. Biasini E, Turnbaugh JA, Unterberger U, Harris DA: Prion protein at the crossroads of physiology and disease. Trends Neurosci 2012, 35(2):92–103.

25. Rambold AS, Müller V, Ron U, Ben-Tal N, Winklhofer KF, Tatzelt J: Stress-protective signalling of prion protein is corrupted by scrapie prions. The EMBO Journal 2008, 27(14):1974–1984.

26. Carroll JA, Chesebro B: Neuroinflammation, Microglia, and Cell-Association during Prion Disease. Viruses 2019, 11(1).

27. Aguzzi A, Zhu C: Microglia in prion diseases. J Clin Invest 2017, 127(9):3230–3239.

28. Mabbott NA, Bradford BM, Pal R, Young R, Donaldson DS: The Effects of Immune System Modulation on Prion Disease Susceptibility and Pathogenesis. Int J Mol Sci 2020, 21(19).

29. Bradford BM, McGuire LI, Hume DA, Pridans C, Mabbott NA: Complete microglia deficiency accelerates prion disease without enhancing CNS prion accumulation. bioRxiv 2021:2021.2001.2005.425436.

30. Carroll JA, Race B, Williams K, Striebel J, Chesebro B: Microglia Are Critical in Host Defense against Prion Disease. J Virol 2018, 92(15):e00549–00518.

31. Carroll JA, Race B, Williams K, Striebel J, Chesebro B: RNA-seq and network analysis reveal unique glial gene expression signatures during prion infection. Mol Brain 2020, 13(1):71.

32. Zhu C, Herrmann US, Falsig J, Abakumova I, Nuvolone M, Schwarz P, Frauenknecht K, Rushing EJ, Aguzzi A: A neuroprotective role for microglia in prion diseases. J Exp Med 2016, 213(6):1047–1059.

33. Kranich J, Krautler NJ, Falsig J, Ballmer B, Li S, Hutter G, Schwarz P, Moos R, Julius C, Miele G et al: Engulfment of cerebral apoptotic bodies controls the course of prion disease in a mouse strain-dependent manner. J Exp Med 2010, 207(10):2271–2281.

34. Gomez-Nicola D, Fransen NL, Suzzi S, Perry VH: Regulation of microglial proliferation during chronic neurodegeneration. J Neurosci 2013, 33(6):2481–2493.

35. Nakagaki T, Ishibashi D, Mori T, Miyazaki Y, Takatsuki H, Tange H, Taguchi Y, Satoh K, Atarashi R, Nishida N: Administration of FK506 from Late Stage of Disease Prolongs Survival of Human Prion-Inoculated Mice. Neurotherapeutics 2020, 17(4):1850–1860.

36. Nazmi A, Field RH, Griffin EW, Haugh O, Hennessy E, Cox D, Reis R, Tortorelli L, Murray CL, Lopez-Rodriguez AB et al: Chronic neurodegeneration induces type I interferon synthesis via STING, shaping microglial phenotype and accelerating disease progression. Glia 2019, 67(7):1254–1276.

37. Makarava N, Kovacs GG, Savtchenko R, Alexeeva I, Budka H, Rohwer RG, Baskakov IV: Stabilization of a prion strain of synthetic origin requires multiple serial passages. J Biol Chem 2012, 287(36):30205–30214.

38. Su EJ, Cao C, Fredriksson L, Nilsson I, Stefanitsch C, Stevenson TK, Zhao J, Ragsdale M, Sun YY, Yepes M et al: Microglial-mediated PDGF-CC activation increases cerebrovascular permeability during ischemic stroke. Acta Neuropathol 2017, 134(4):585–604.

39. Makarava N, Mychko O, Molesworth K, Chang JC, Henry RJ, Tsymbalyuk N, Gerzanich V, Simard JM, Loane DJ, Baskakov IV: Region-Specific Homeostatic Identity of Astrocytes Is Essential for Defining Their Response to Pathological Insults. Cells 2023, 12(17).

40. Franceschini A, Strammiello R, Capellari S, Giese A, Parchi P: Regional pattern of microgliosis in sporadic Creutzfeldt-Jakob disease in relation to phenotypic variants and disease progression. Neuropathol Appl Neurobiol 2018, 44(6):574–589.

41. Makarava N, Chang JC-Y, Molesworth K, Baskakov IV: Region-specific glial homeostatic signature in prion diseases is replaced by a uniform neuroinflammation signature, common for brain regions and prion strains with different cell tropism. Neurobiology of Disease 2020, 137(1):e104783.

42. VanRyzin JW, Marquardt AE, Argue KJ, Vecchiarelli HA, Ashton SE, Arambula SE, Hill MN, McCarthy MM: Microglial Phagocytosis of Newborn Cells Is Induced by Endocannabinoids and Sculpts Sex Differences in Juvenile Rat Social Play. Neuron 2019, 102(2):435–449.e436.

43. Barcia C, Ros CM, Annese V, Carrillo-de Sauvage MA, Ros-Bernal F, Gómez A, Yuste JE, Campuzano CM, de Pablos V, Fernandez-Villalba E et al: ROCK/Cdc42-mediated microglial motility and gliapse formation lead to phagocytosis of degenerating dopaminergic neurons in vivo. Sci Rep 2012, 2:809.

44. Carroll JA, Striebel JF, Rangel A, Woods T, Phillips K, Peterson KE, Race B, Chesebro B: Prion Strain Differences in Accumulation of PrPSc on Neurons and Glia Are Associated with Similar Expression Profiles of Neuroinflammatory Genes: Comparison of Three Prion Strains. PLoS Pathog 2016, 12(4):e1005551.

45. Nemes-Baran AD, White DR, DeSilva TM: Fractalkine-Dependent Microglial Pruning of Viable Oligodendrocyte Progenitor Cells Regulates Myelination. Cell Rep 2020, 32(7):108047.

46. Ferrer I, Casas R, Rivera R: Parvalbumin-immunoreactive cortical neurons in Creutzfeldt-Jakob disease. AnnNeurol 1993, 34:864–866.

47. Guentchev M, Groschup MH, Kordek R, Liberski PP, Budka H: Severe, Early and Selective Loss of a Subpopulation of GABAergic Inhibitory Neurons in Experimental Transmissible Spongiform Encephalopathies. Brain Pathology 1998, 8(4):615–623.

48. Guentchev M, Hainfellner J-A, Trabattoni GR, Budka H: Distribution of Parvalbumin-Immunoreactive Neurons in Brain Correlates with Hippocampal and Temporal Cortical Pathology in Creutzfeldt-Jakob Disease. Journal of Neuropathology & Experimental Neurology 1997, 56(10):1119–1124.

49. Franklin SL, Love S, Greene JR, Betmouni S: Loss of Perineuronal Net in ME7 Prion Disease. J Neuropathol Exp Neurol 2008, 67(3):189–199.

50. Kavanagh E, Rodhe J, Burguillos MA, Venero JL, Joseph B: Regulation of caspase-3 processing by cIAP2 controls the switch between pro-inflammatory activation and cell death in microglia. Cell Death & Disease 2014, 5(12):e1565–e1565.

51. Burguillos MA, Deierborg T, Kavanagh E, Persson A, Hajji N, Garcia-Quintanilla A, Cano J, Brundin P, Englund E, Venero JL et al: Caspase signalling controls microglia activation and neurotoxicity. Nature 2011, 472(7343):319–324.

52. Wakselman S, Béchade C, Roumier A, Bernard D, Triller A, Bessis A: Developmental Neuronal Death in Hippocampus Requires the Microglial CD11b Integrin and DAP12 Immunoreceptor. The Journal of Neuroscience 2008, 28(32):8138.

53. Allendorf DH, Puigdellívol M, Brown GC: Activated microglia desialylate their surface, stimulating complement receptor 3-mediated phagocytosis of neurons. Glia 2020, 68(5):989–998.

54. Parchi P, Giese A, Capellari S, Brown P, Schulz-Schaeffer W, Windl O, Zerr I, Budka H, Kopp N, Piccardo P et al: Classification of sporadic Creutzfeldt-Jakob disease based on molecular and phenotypic analysis of 300 subjects. Ann Neurol 1999, 46:224–233.

55. Liu B, Wang K, Gao HM, Mandavilli B, Wang JY, Hong JS: Molecular consequences of activated microglia in the brain: overactivation induces apoptosis. J Neurochem 2001, 77(1):182–189.

56. Sinha A, Kushwaha R, Molesworth K, Mychko O, Makarava N, Baskakov IV: Phagocytic Activities of Reactive Microglia and Astrocytes Associated with Prion Diseases Are Dysregulated in Opposite Directions. Cells 2021, 10(7):1728.

57. Cserép C, Pósfai B, Lénárt N, Fekete R, László ZI, Lele Z, Orsolits B, Molnár G, Heindl S, Schwarcz AD et al: Microglia monitor and protect neuronal function through specialized somatic purinergic junctions. Science 2020, 367(6477):528–537.

58. Makarava N, Chang JC-Y, Molesworth K, Baskakov IV: Posttranslational modifications define course of prion strain adaptation and disease phenotype. The Journal of Clinical Investigation 2020, 130(8):4382–4395.

59. Srivastava S, Katorcha E, Makarava N, Barrett JP, Loane DJ, Baskakov IV: Inflammatory response of microglia to prions is controlled by sialylation of PrP^Sc^. Sci Rep 2018, 8(1):e11326.

60. Scheckel C, Imeri M, Schwarz P, Aguzzi A: Ribosomal profiling during prion disease uncovers progressive translational derangement in glia but not in neurons. Elife 2020, 9.

61. Lui H, Zhang J, Makinson Stefanie R, Cahill Michelle K, Kelley Kevin W, Huang H-Y, Shang Y, Oldham Michael C, Martens Lauren H, Gao F et al: Progranulin Deficiency Promotes Circuit-Specific Synaptic Pruning by Microglia via Complement Activation. Cell 2016, 165(4):921–935.

62. Hansen DV, Hanson JE, Sheng M: Microglia in Alzheimer’s disease. The Journal of Cell Biology 2018, 217(2):459–472.

63. Ugalde CL, Lewis V, Stehmann C, McLean CA, Lawson VA, Collins SJ, Hill AF: Markers of A1 astrocytes stratify to molecular sub-types in sporadic Creutzfeldt-Jakob disease brain. Brain Communications 2020, 2(2):fcaa029.

64. Baskakov IV, Katorcha E: Multifaceted role of sialylation in prion diseases. Front Neurosci 2016, 10(1):e358.

65. Katorcha E, Makarava N, Savtchenko R, Baskakov IV: Sialylation of the prion protein glycans controls prion replication rate and glycoform ratio. Sci Rep 2015, 5(1):16912.

66. Baskakov IV, Katorcha E, Makarava N: Prion Strain-Specific Structure and Pathology: A View from the Perspective of Glycobiology. Viruses 2018, 10(12):723.

67. Makarava N, Chang JC-Y, Baskakov IV: Region-Specific Sialylation Pattern of Prion Strains Provides Novel Insight into Prion Neurotropism. International Journal of Molecular Sciences 2020, 21(3):828.

68. Sandberg MK, Al-Doujaily H, Sharps B, De Oliveira MW, Schmidt C, Richard-Londt A, Lyall S, Linehan JM, Brandner S, Wadsworth JD et al: Prion neuropathology follows the accumulation of alternate prion protein isoforms after infective titre has peaked. Nat Commun 2014, 5:e4347.

69. Makarava N, Mychko O, Chang JC-Y, Molesworth K, Baskakov IV: The degree of astrocyte activation is predictive of the incubation time to prion disease. Acta Neuropathologica Communications 2021, 9(1):87.

70. Cunningham CL, Martínez-Cerdeño V, Noctor SC: Microglia regulate the number of neural precursor cells in the developing cerebral cortex. J Neurosci 2013, 33(10):4216–4233.

71. Zhao L, Zabel MK, Wang X, Ma W, Shah P, Fariss RN, Qian H, Parkhurst CN, Gan WB, Wong WT: Microglial phagocytosis of living photoreceptors contributes to inherited retinal degeneration. EMBO Mol Med 2015, 7(9):1179–1197.

72. Puigdellívol M, Milde S, Vilalta A, Cockram TOJ, Allendorf DH, Lee JY, Dundee JM, Pampuščenko K, Borutaite V, Nuthall HN et al: The microglial P2Y(6) receptor mediates neuronal loss and memory deficits in neurodegeneration. Cell Rep 2021, 37(13):110148.

