## Supplementary figures and images for "Engulfment of viable neurons by reactive microglia in prion diseases"

### Fig. S1

Fig. S1

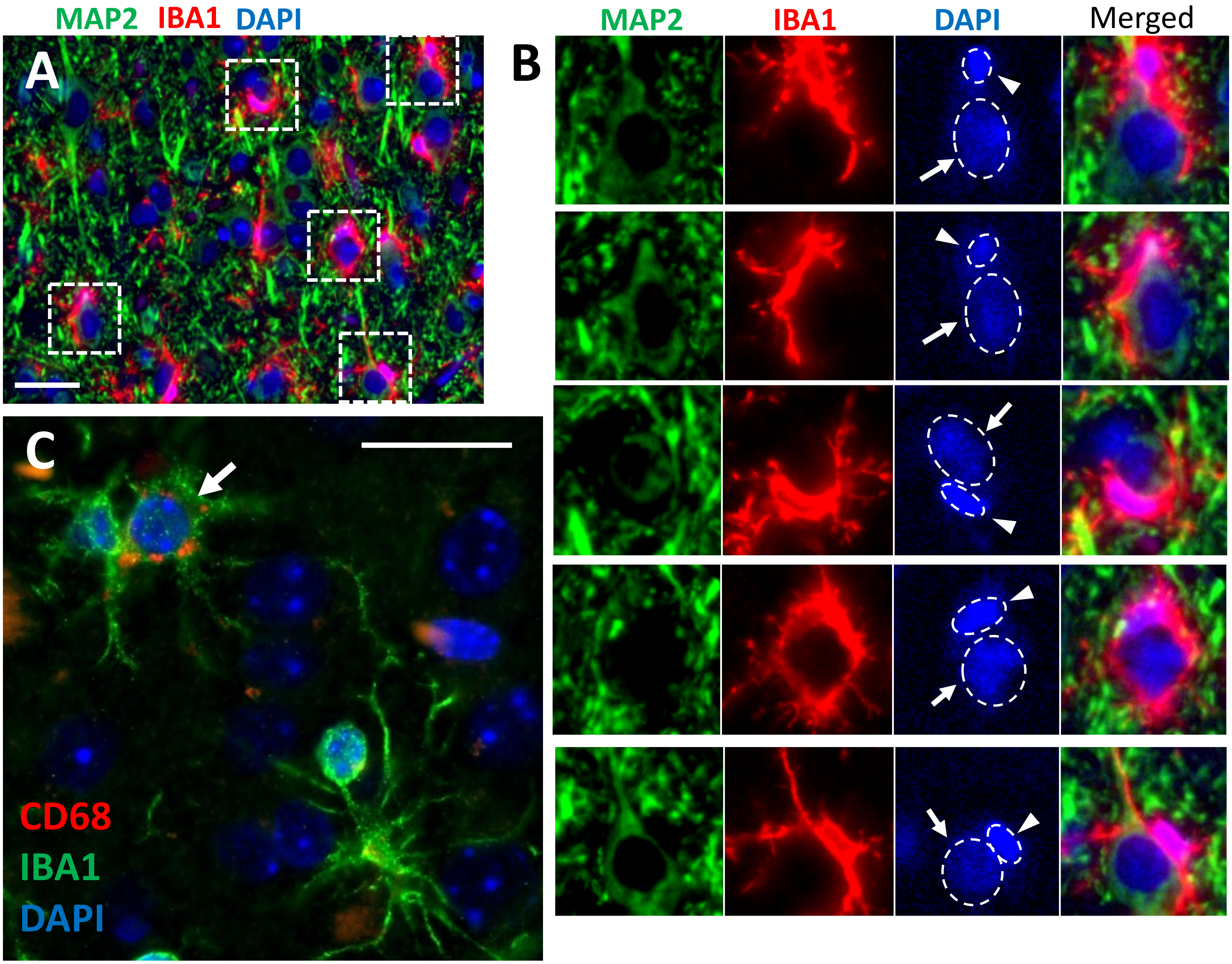

### Fig. S2

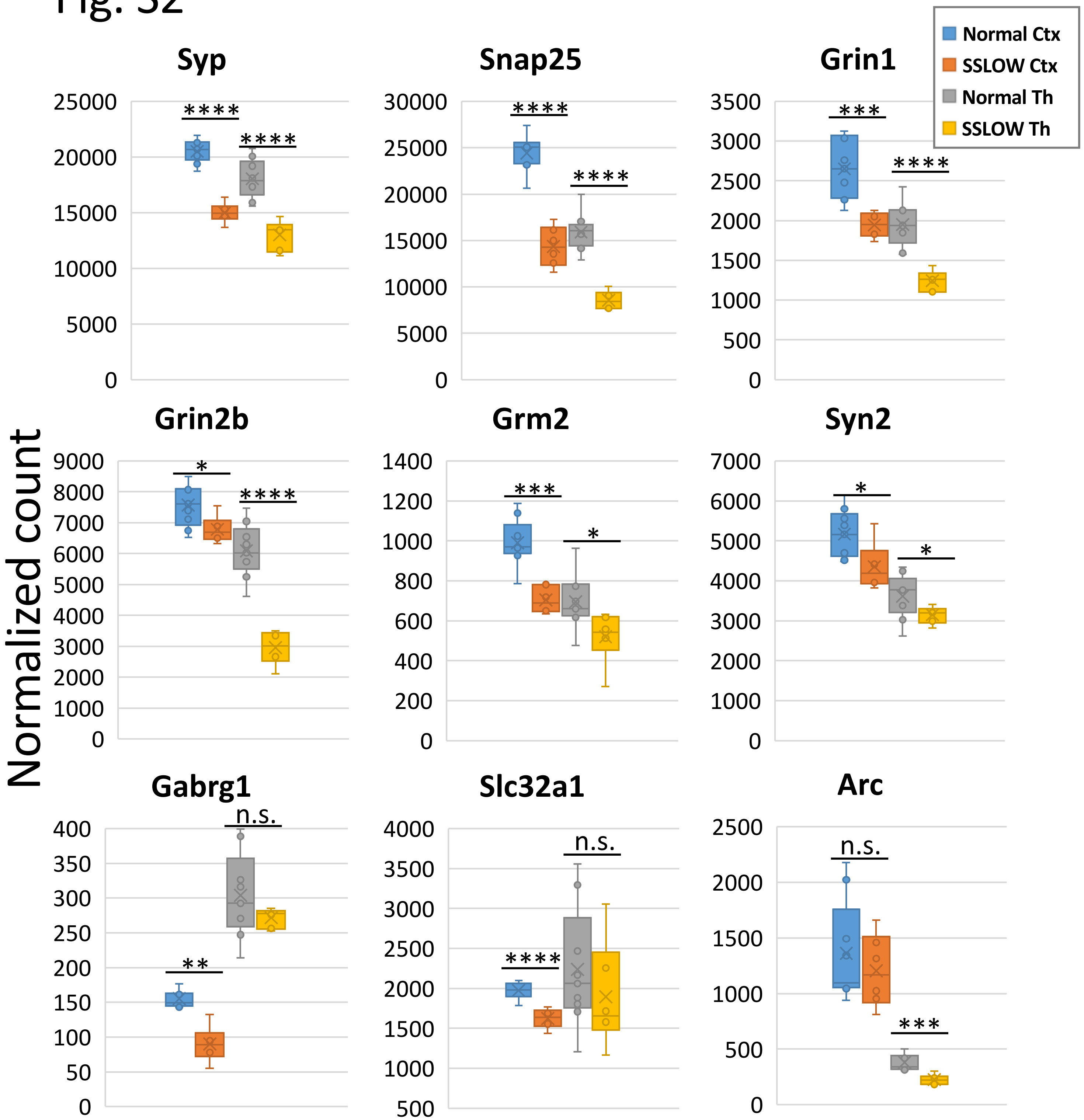

### Fig. S3

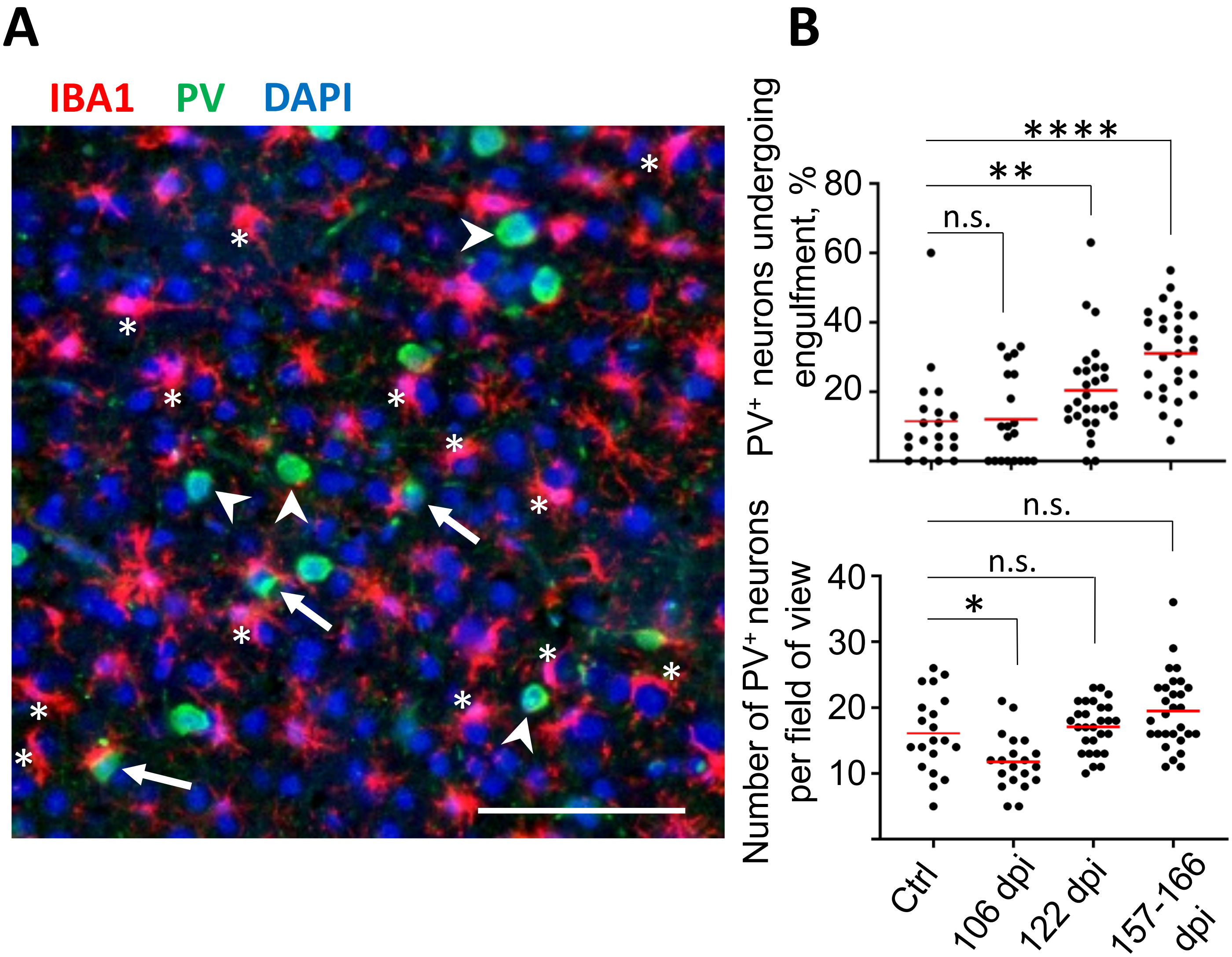

### Fig. S4

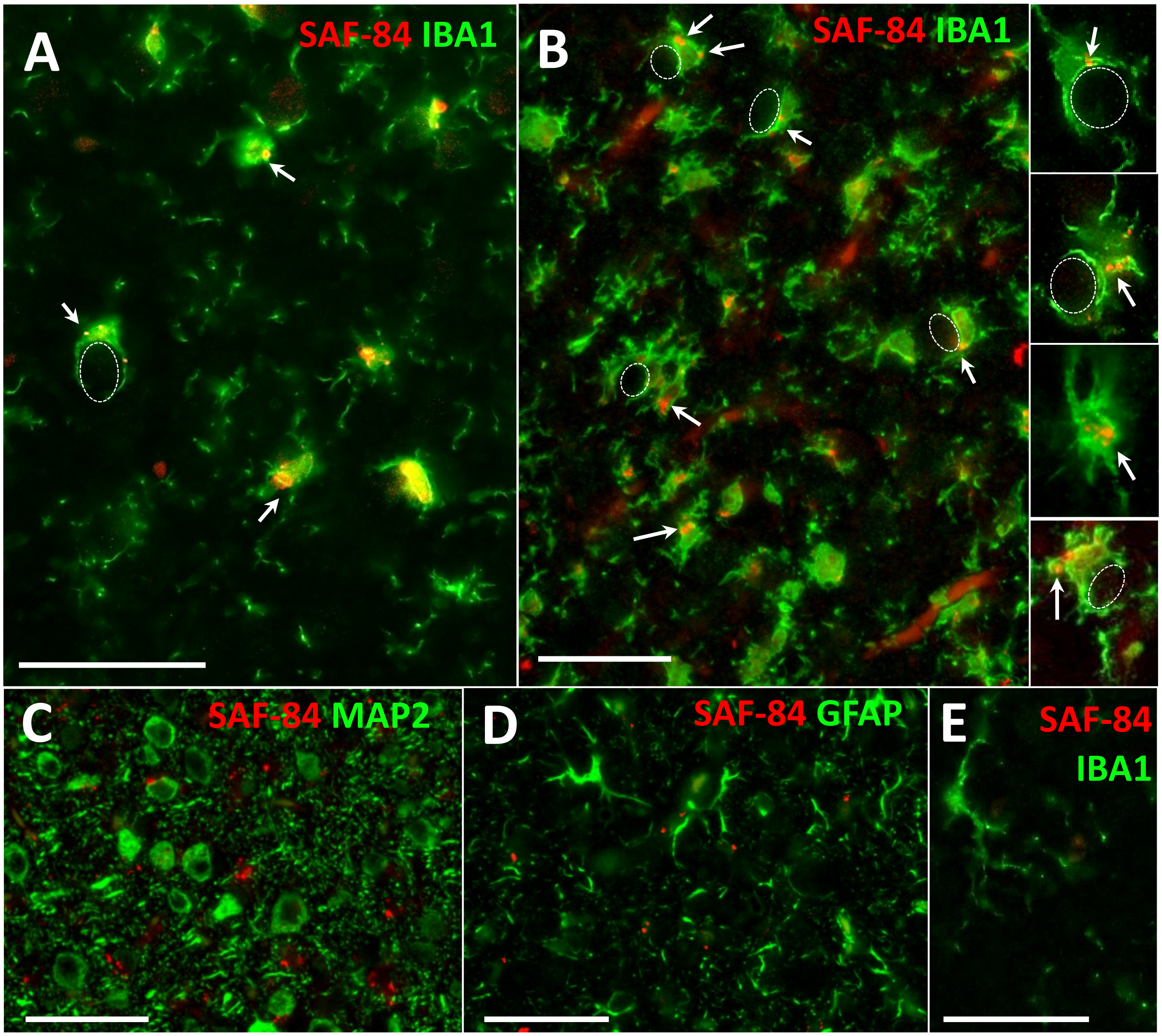

### Fig. S5

#### Slide 1
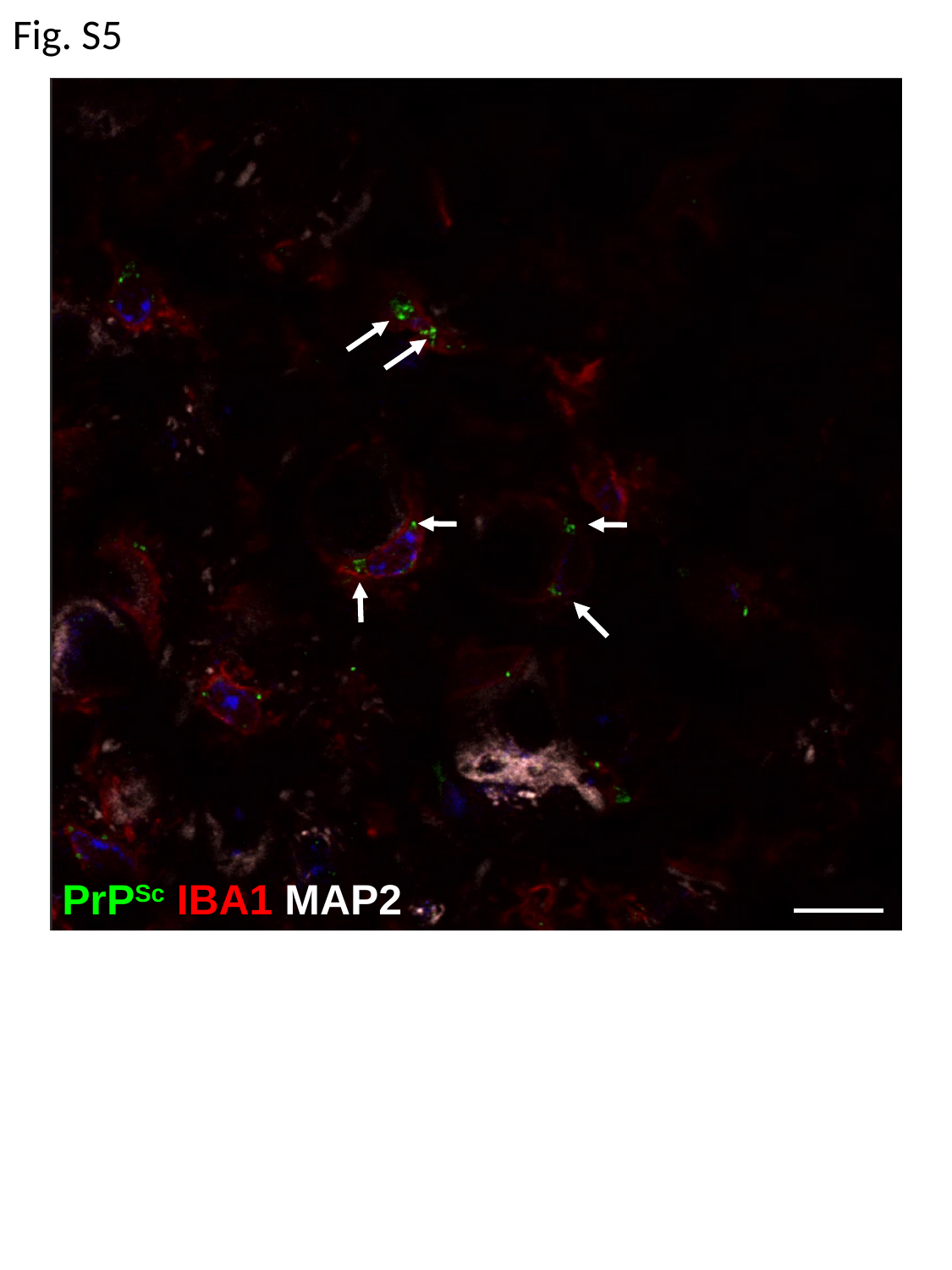

Fig. S5
PrP^Sc^ IBA1 MAP2

### Fig. S6

Fig. S6

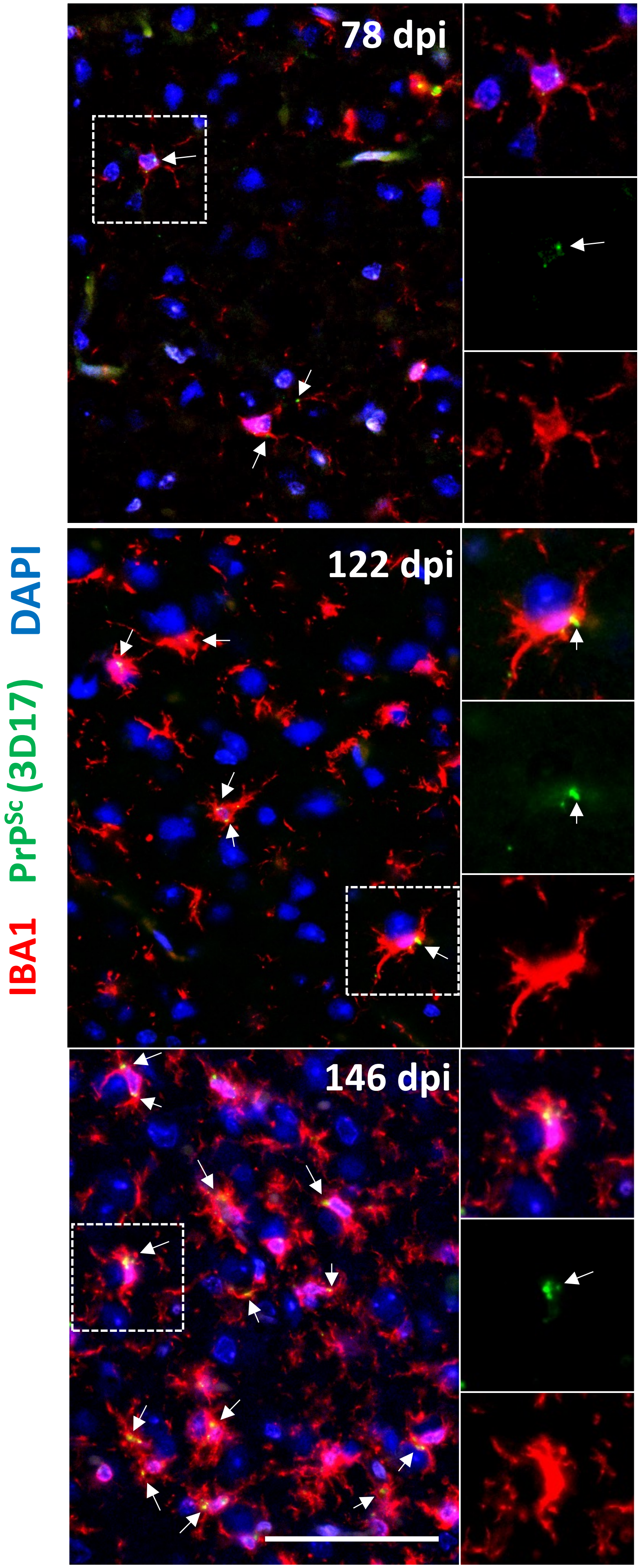
